# Neutrophils respond with pathogen-specific defenses during bacterial pneumonia

**DOI:** 10.1101/2025.04.17.649365

**Authors:** Riley M.F. Pihl, Kevyn R. Martins, Yewoo Lee, Lilly Patneaude, Lee J. Quinton, Joseph P. Mizgerd, Anna C. Belkina, Katrina E. Traber

## Abstract

Neutrophils have historically been envisioned as a homogenous population of short-lived innate immune cells that migrate to sites of infection, kill pathogens, and die. Recent work, including studies in pneumonia models, has shown that neutrophil transcriptomes reflect the environment from which they were isolated. We used high-parameter spectral flow cytometry to compare and contrast a wide array of surface proteins on neutrophils from different tissues, infections, host age, pathogen virulence, and across multiple time-points of pneumonia. Circulating and airspace neutrophils consistently differed, and surface protein phenotypes unique to each infection setting were identified, revealing tissue-specific and microbe-specific neutrophil plasticity. Phenotypic differences in circulating neutrophils from mice infected with different pathogens (*E. coli*, *S. pneumoniae*, *S. aureus*, and *P. aeruginosa*) identified, even in the absence of bacteremia. Neutrophil activation state was diminished with less virulent pathogens and host age. In the airspace, VISTA, CD200R, and PD-L1 were selectively high on BAL neutrophils (BALN) during *S. pneumoniae* infection, and we identified pro-degranulation-like (CD88^High^ VISTA^High^ PD-L1^+^ CD101^-^) neutrophils in *S. pneumoniae* and pro-phagocytosis-like (CD101^+^ CD18^Low^ PD-L1^-^) neutrophils in *E. coli* infections. Stimulation of VISTA with its ligand VISG-3 enhanced the neutrophil respiratory burst, degranulation, and killing of *S. pneumoniae* but not *E. coli*. We conclude that neutrophil cell surface protein expression depends on anatomic location and infection type, resulting in pathogen-specific neutrophil-mediated immune defense in discreet areas of the pneumonic lung.

**Graphical Abstract:** In brief, Pihl et al. have found that neutrophil cell surface phenotype varies drastically based on tissue, time post-infection, and infection. BALN from early infections have higher activation and maturation statuses, while blood neutrophils are more ‘migration primed,’ and late infections have more immune-suppressive and altered pathogen killing statuses. Neutrophil phenotype is skewed towards pro-phagocytosis associated marker expression on BALN from *E. coli*-infected mice while *S. pneumoniae* results in a pro-degranulation phenotype. In vitro BMN stimulation of VISTA with VSIG-3 results in degranulation, respiratory burst, and pathogen specific killing of *S. pneumoniae* but not *E. coli*.

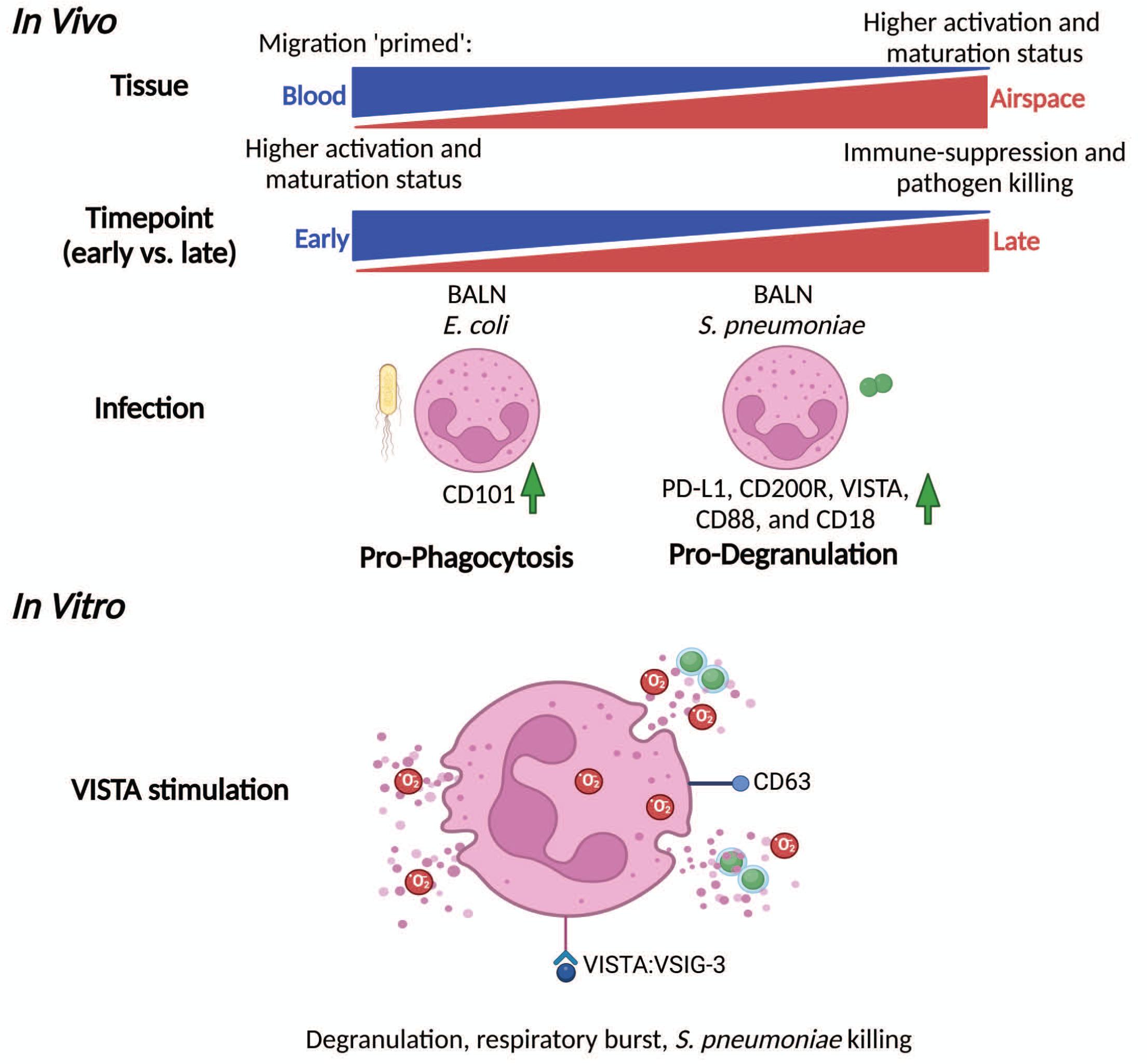

## Introduction

Respiratory infections remain a prominent cause of global death and morbidity, despite pathogen-specific therapeutics which include antibiotics and vaccines effective against common respiratory pathogens such as *S. pneumoniae* [1]. High levels of inflammatory cytokines and an unchecked innate immune response are hallmarks of the immune dysregulation that drive disease in pneumonia, leading to tissue damage and increased patient morbidity and mortality [2, 3].

Developing novel host-directed therapeutics to correct immune dysfunction is a current priority of pneumonia research [4], demanding a better understanding of signals controlling immune dysregulation [3, 5].

Neutrophils are short-lived innate immune cells that respond quickly in infection to clear pathogens yet can also generate bystander tissue damage via inflammatory cytokine production and other effector functions. If unchecked, these neutrophil actions can lead to severe lung injury, acute respiratory distress syndrome (ARDS) and/or sepsis [6, 7], yet patients with neutropenia, i.e. insufficient neutrophil response, are at risk of severe bacterial infections [8]. Therefore, neutrophil responses require strict calibration, as both too mild or too aggressive of a response can drive disease.

Neutrophils are produced in the bone marrow, circulate in the blood, and then migrate to sites of infection, where they kill pathogens via several discrete processes; degranulation that releases reactive oxygen species (ROS) and bactericidal enzymes such as myeloperoxidase (MPO) and neutrophil elastase (NE), phagocytosis, and production of neutrophil extracellular traps (NETs) [9]. While previously thought to be solely a pathogen killing effector cell type, recent work has shown that neutrophils can fulfill many additional functions, which include but were not limited to regulation of lymphocytes [10, 11], tumor propagation or elimination [12], and resolution of inflammation [13].

Further recent work, including our own, has shown that neutrophils are highly transcriptionally dynamic and influenced by their microenvironment. The neutrophil transcriptome is distinct between neutrophils isolated from different disease models and from different tissues within a given model [14–17]. Our own recent work has shown that thousands of genes were significantly changed between neutrophils isolated from the circulation or bronchioloalveolar lavage (BAL) fluid, and between vascular, interstitial and alveolar sites [18]. BAL neutrophil heterogeneity was also observed, where an “early” or naïve-like transcriptome appears to mature into two distinct “late” or mature-like phenotypes. The current work builds on prior transcriptomic characterizations by examining neutrophil cell surface protein expression in multiple experimental models of pneumonia.

## Results

### Neutrophil phenotype stratifies predominantly by tissue compartment and infection type

To investigate neutrophil phenotypes across multiple models of pneumonia, we developed a 24-parameter full spectrum neutrophil flow cytometry panel, choosing proteins of interest from our own transcriptomic data as well as from the wider body of neutrophil data. For a listing of antibody source information, aliases, and putative functions, please see **Supplemental Table 1**. Our pneumonia models (overview **Figure 1A**) utilized severe lethal strains of *E. coli* (EC) or *S. pneumoniae* (serotype 3, SP3), or mild/non-lethal *S. pneumoniae* (serotype 19f, SP19f). In our experience, EC-infected mice are moribund at 48h, where SP3-infected mice are moribund by 72-hours. We therefore defined early pneumonia as 6-hour EC infection and 24-hour SP3 infection, versus late pneumonia as EC 24-hours and SP3 48-hours. Finally, we explored neutrophils from aged mice infected with SP3 for 24h-hours. Our gating strategy is shown in **Supplemental Figure 1A** and example channels demonstrating lack of batch-effects in **Supplemental Figure 1B-G**. In **Figure 1** we present an overview of the data from these studies. In **Figure 1B** is a dimensionality reduction opt-SNE plot depicting neutrophils from all experimental conditions, colored by the thirty-five PhenoGraph clusters determined in our initial analysis and separate opt-SNE plots from circulating or BAL-derived neutrophils. When neutrophils from specific experimental groups were plotted on their own opt-SNE maps, clusters separate well by experimental groups (**Figure 1CD**) and these differences will be discussed in following sections. Resident airway neutrophils are typically not observed during homeostasis and were thus not collected. Example markers that distinguish circulating blood neutrophils (CN, **Figure 1E**) and BAL neutrophils (BALN, **Figure 1F**) are presented as pseudocolor overlays (all other overlays: **Supplemental Figure 1H**). In **Figure 1G** is a graph demonstrating the relative contribution of surface protein expression (heatmap), and experimental group (bar graph) to the identified PhenoGraph clusters (statistical analysis of experimental group contributions: **Supplemental Table 2**). Columns are hierarchically ordered by X, and rows by Y. PhenoGraph clusters appear to hierarchically group by tissue and pathogen, where clusters enriched in CN from uninfected and EC-infected (EC-CN) conditions group towards the top of the heatmap (P30, P23, P24, P28, P8, P35, P20, P15, and P10), followed by CN from SPN-infected (SPN- CN; P7, P12, P3, P5, P14, and P29), BALN from SPN-infected (SPN-BALN; P1, P2, P22, P6, and P13) and lastly BALN from EC-infected (EC-BALN; P11, P4, P21, P9, P26, and P16) (**Figure 1G**). Further, some clusters are enriched in neutrophils from early (P10, P15, P3, P5, P11, P21, and P1), late, (P18, P20, P12, P17, P4, P16, and P6), mild (P7 and P1), or aged mouse (P14 and P19) conditions (**Figure 1G**). This suggests that tissue type and infection are two of the driving variables behind observed differences in neutrophil phenotypes in this dataset, though timepoint, age, and virulence can also have an impact. PhenoGraph clusters were investigated for sex-based differences, though none were found (**Supplemental Figure 1I**). Blood CFU and lung proteinaceous edema (as a measure of lung damage) were quantified for each subject (**Extended Data Figure 1AB**) and clusters that were enriched in mice with bacteremia or proteinaceous edema were also identified and discussed in later sections (**Extended Data Figure 1CD**). Overall, neutrophil phenotype was highly tuned by tissue and infection model, and differences were also observed with timepoint, age, and lethality.

**Figure 1.**
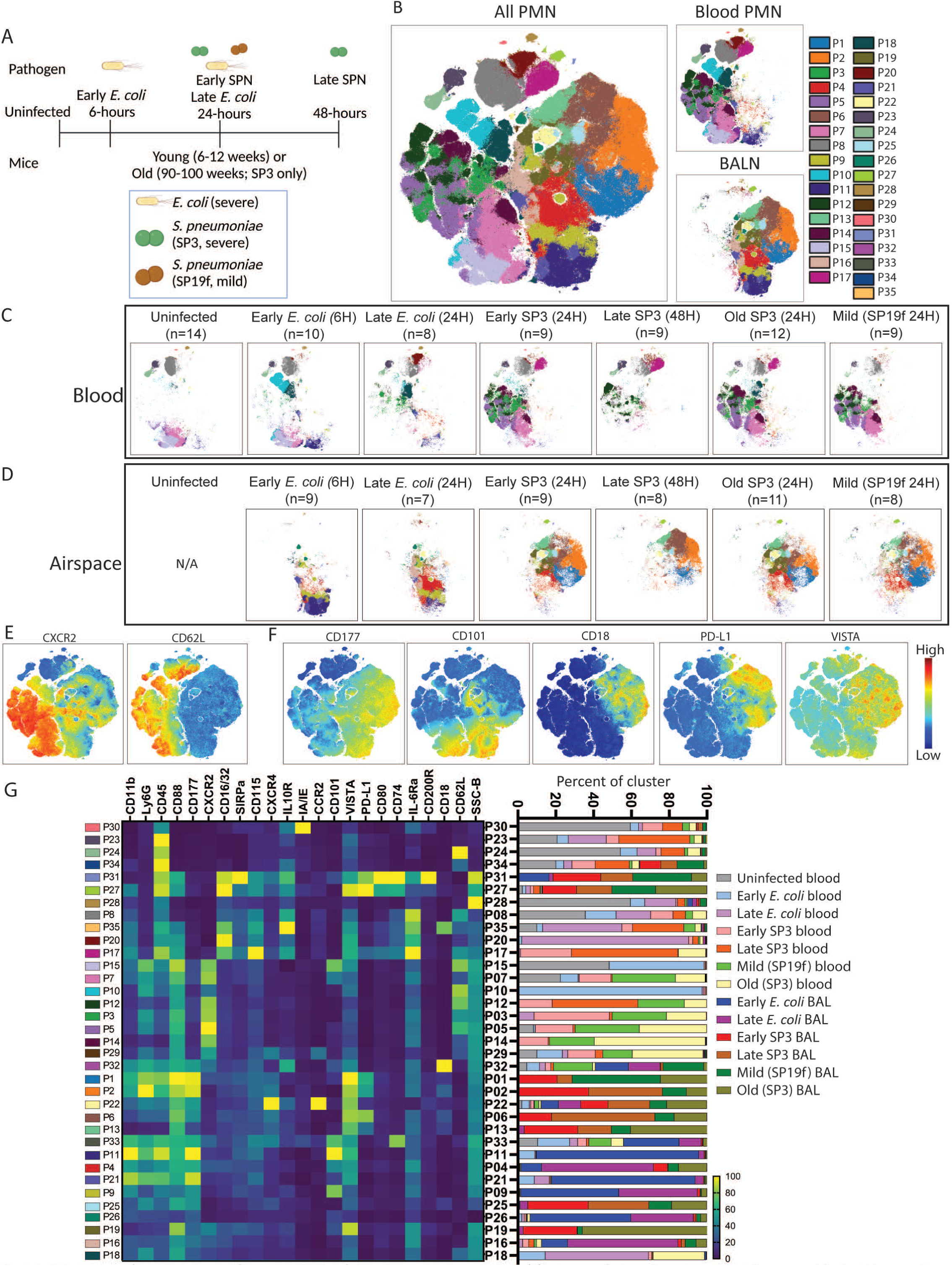
High-parameter flow cytometry stratifies neutrophils by infection, time-point, tissue, and age. (**A**) Schematic of all samples and experimental groups used for the high-parameter flow cytometry dataset. Blood neutrophils were collected from uninfected mice, and blood and BAL neutrophils were collected from mice infected with EC for 6-, EC for 24-, SP3 for 24-, SP3 in aged mice for 24-, SP19f for 24-, and SP3 for 48-hours. (**B**) Cells from all experimental groups was plotted on an opt-SNE map with PhenoGraph clusters overlayed by color, with secondary plots depicting blood neutrophils or BAL neutrophils only. Subsets of the opt-SNE plot were also plotted to show where cells from both (**C**) blood and (**D**) BAL experimental groups were located on the map, and to which clusters they belonged. Pseudocolor overlays of features that distinguish (**E**) blood and (**F**) BAL neutrophils were plotted. (**G**) PhenoGraph clusters were hierarchically organized and plotted in a heatmap with each parameter on the X-axis (normalized to unstained blood neutrophils) and the PhenoGraph clusters on the Y-axis which were hierarchically grouped based on the MFls of each cluster. Stacked bar graphs to the right show the fraction of each experimental group represented in each cluster.

### Early circulating neutrophils are more ‘migration primed’ than late circulating neutrophils

As noted above, subsets of PhenoGraph clusters were associated with neutrophils from certain experimental conditions. We examined the surface marker profiles associated with these experimental conditions, beginning with clusters associated with CNs (**Figure 2**). Clusters P15 and P10 were associated with early EC-CN and have higher CD62L (L-selectin) and CXCR2 compared to late EC clusters (P18 and P20), suggesting that early circulating neutrophils were “migration primed” while later circulating neutrophils were less prepared to migrate (**Figure 2AB**). Among SPN-CN-associated clusters, both CD62L and CXCR2 are elevated early and only a subset (P17) have a late decrease in CXCR2. Interestingly, CD101 appears to be strongly enriched on early EC-CN cluster P15, and to a lesser extent on early SPN-CN cluster P5. CD101 has been used as a neutrophil maturation marker where CD101^+^ neutrophils are more mature than CD101^-^, and recent studies found that CD101^+^ neutrophils phagocytose more bacteria and do so faster than CD101^-^ neutrophils [19–21]. Interestingly, P10 (enriched in early EC-CN) and P3 (enriched in early SPN-CN), do not have increased CD101, suggesting that there is a mix of circulating neutrophils in early infections that either are or are not pro-phagocytosis. Further, both late clusters P20 (EC) and P17 (SP3) are CD16/32^High^ CD177^Low^ SIRPα^+^ Ly6G^Low^.

**Figure 2.**
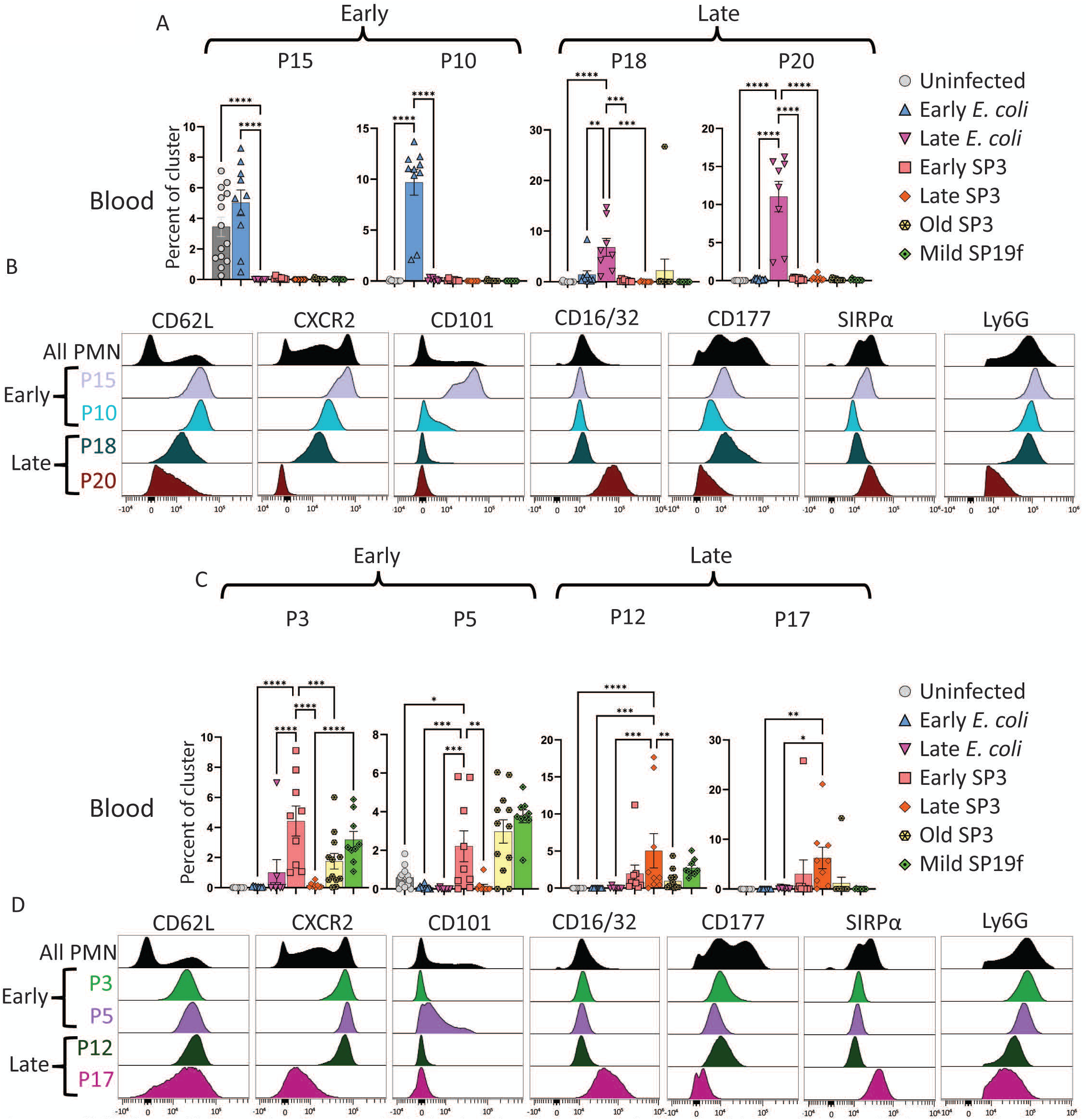
Specific blood neutrophil clusters and markers are associated with early and late infections. Bar graphs show selected clusters with a significantly higher fraction of blood neutrophils from mice infected with (**A**) uninfected or EC or (**C**) SPN. Histograms show specific marker expression of each (**B**) EC, (**D**) or SPN clusters. Statistics used were one-way ANOVAs with Bonferroni multiple comparisons corrections where * = P < 0.05, ** = P < 0.01, *** = P < 0.001, and **** = P < 0.0001. Error bars show standard error of the mean (SEM).

Interestingly, CD16^High^ neutrophils have been previously reported to clear blood infections via phagocytosis when serum IgG concentration is high [22], while SIRPα is a “don’t eat me” signal used by host cells and bacteria to prevent phagocytosis [23–25]. Additionally, P17, P18, and P20 are enriched in mice with bacteremia (**Extended Data Figure 1C**), suggesting that these groups of neutrophils circulating late in infection that have had direct pathogen exposure may be pro- bacteria killing. The presence of SIRPα on CD16^High^ neutrophils may have the additional function of preventing improper targeting of neutrophils that otherwise would be primed for killing by antibody-dependent cellular cytotoxicity (ADCC).

### Circulating neutrophils differ by infection type, with the most pronounced differences occurring during bacteremia

A striking feature of our circulating neutrophil dataset was that not only can we identify EC and SP3 associated circulating neutrophil phenotypes, but pathogen-associated CN subsets exist even in settings without bacteremia (EC cluster P10 **Figure 2A**; SP19f clusters P3 and P5 **Figure 2C**), suggesting that circulating factors, rather than direct pathogen interaction may drive neutrophil phenotype. We performed a second set of pneumonia models, across four bacteria species (SP3, EC, *P. aeruginosa* and *S. aureus*) at levels not expected to develop bacteremia (low dose, LD).

We evaluated circulating neutrophils from these infections by spectral flow, utilizing a subset of markers from our initial dataset (**Extended Data Figure 2A-C**). Each infection group has a discrete cluster and marker expression pattern that predominantly separate by CXCR2, CD62L, SIRPα, CD66a, and SSC expression, suggesting that neutrophil phenotype is tuned in the blood while *en route* to sites of infection and before direct pathogen exposure. To directly compare this new dataset with our original, compatible channels were normalized in OMIQ.ai using CytoNorm [26] (**Extended Data Figure 3**). In this new combined circulating neutrophil dataset, severe EC or SP3 (early and late) are highly distinct from their low dose counterparts or the low dose *S. aureus* or *P. aeruginosa* (**Extended Data Figure 3BC**). Circulating neutrophils from severe EC infections have higher CD62L and lower CXCR2 relative to low dose EC (**Extended Data Figure 3CD**). Neutrophils from severe SP3 are the only experimental group with high PD- L1 expression, and both lethal infection groups have less CD101 expression relative to uninfected or low dose infection groups (**Extended Data Figure 3CD**). Taken together, these data strongly suggests that neutrophils begin divergent pathogen-specific trajectories even without direct pathogen exposure, though bacteremia drives more dramatic changes.

### Early EC-BALN are more activated than late, though both EC-BALN timepoints retain high pro-phagocytosis CD101 expression phenotypes

As noted above, clusters predominantly composed of BALN group separately from CN-associated clusters. Furthermore, clusters composed primarily of EC-infected BALN (EC-BALN - P11, P21, P9, P26, P4, and P16) largely group separately from SPN-infected neutrophils (SPN-BALN - P1, P2, P6, P13, and P19) (**Figure 3AC**). In EC infections, early-BALN (P11 and P21) clusters express high levels of CD11b, CD177, and Ly6G, and are distinguished by CD101 expression (high on P11 but not P21) (**Figure 3AB**). In contrast, clusters P9 and P26, which were enriched in both early and late EC infections (grouped together as “severe” infections), express lower levels of CD11b, CD177 and Ly6G, yet still separate by expression of CD101 (P9 higher than P26) (**Figure 3B**). Lastly, late EC-BALN clusters (P4 and P16) express lower levels of CD11b, Ly6G, and CD177, but higher CXCR2. These clusters also separate based on CD101, (high on P4), additionally P4 has the highest VISTA expression of EC-BALN clusters (**Figure 3AB**). Overall, it appears that EC-BALN neutrophils initially have high expression of activation markers and were replaced over time by more naïve and less primed phenotypes, that may be pro-phagocytosis.

**Figure 3.**
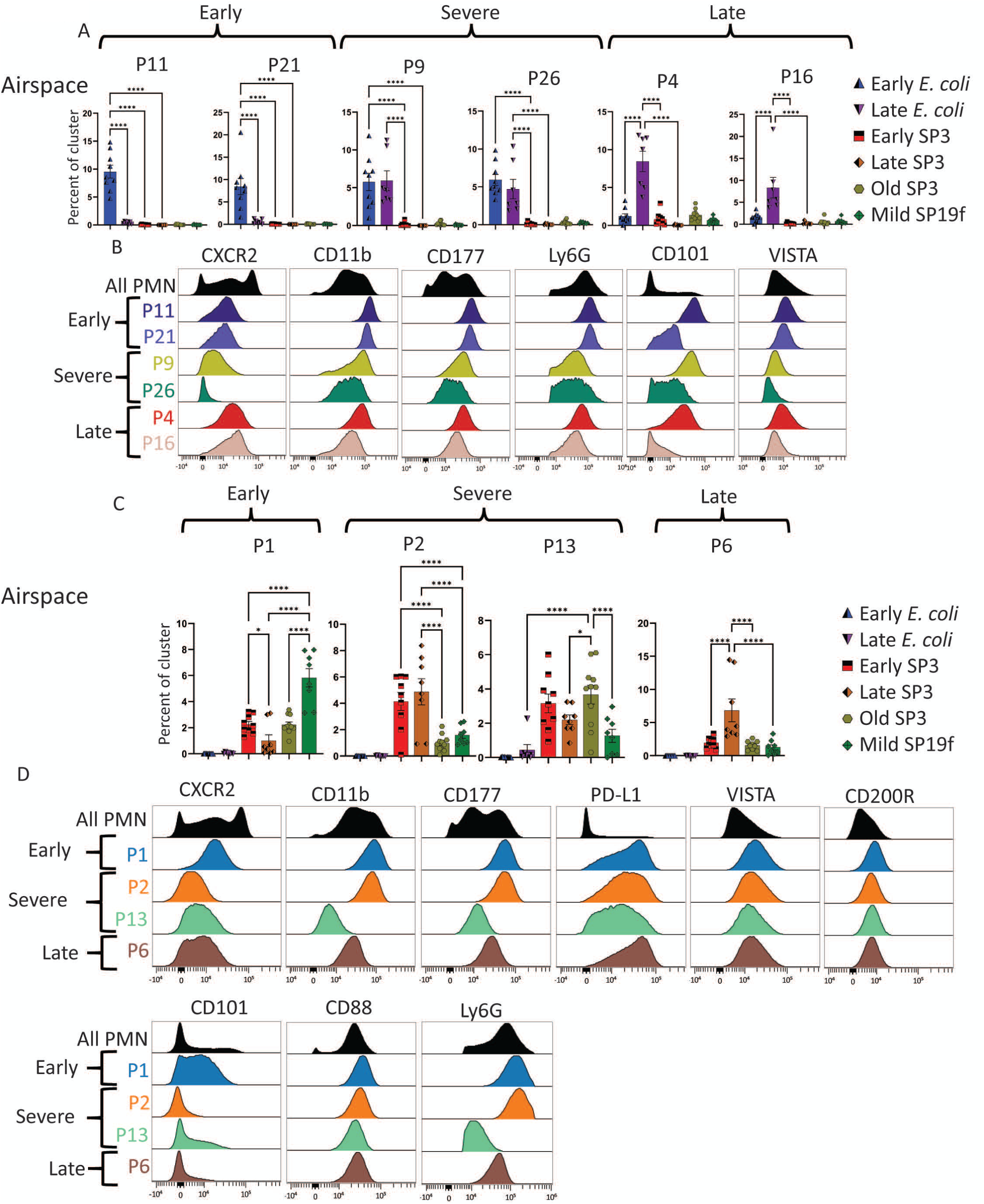
Specific BALN clusters and markers are associated with early and late infections. Bar graphs show selected clusters with a significantly higher fraction of BALNs from mice infected with (**A**) EC or (**C**) SPN. Histograms show specific marker expression of each (**B**) EC, (**D**) or SPN clusters. Statistics used were one-way ANOVAs with Bonferroni multiple comparisons corrections where * = P < 0.05, ** = P < 0.01, *** = P < 0.001, and **** = P < 0.0001. Error bars show standard error of the mean (SEM).

### Immune regulators PD-L1, CD200R, and VISTA were significantly higher on BALNs associated with SPN-infected mice

Compared to EC-BALN clusters, SPN-BALN (P1, P2, P6, P13 and P19) have higher expression of PD-L1, CD200R, VISTA, CD88, and CD18, yet lower on average CD101 (**Figure 1G**; **Figure 3CD)**. SPN-BALN-associated clusters can also be further subset into those associated with early infection (P1), early and late infection (“severe” P2, P13), and late infection (P6). P1 and P2 are phenotypically similar, though P1 expresses higher CD101 and CXCR2 compared to P2 (**Figure 3D**), which is consistent with an early more mature and recently migrated phenotype. Also akin to EC early and late phenotypes, early SPN P1 has higher activation and maturation markers CD11b, CD177, and CD101 compared with late-SP3 associated P6 (**Figure 3CD**). Unique to SPN BALNs, early SPN BALN (P1) appear to have modestly higher VISTA and CD200R but lower PD-L1 compared to late (P6) neutrophils. CD88 (C5aR) expression is highest on early SP3-BALNs (**Figure 3D**), and activation is known to result in chemotaxis and degranulation [27, 28]. Taken together, these data suggest that early versus late neutrophils have some shared characteristics across infections such as lower activation states, while different functions such as phagocytosis and degranulation across time and infections may be pathogen-specific.

### Mild infections and advanced age alter the activation and immunosuppressive state of neutrophils

In addition to looking at the effect of pathogen type on neutrophil function, we also investigated the effect of age (6-12 week versus 90week) as well as pathogen severity (SP3 versus SP19). Both virulence and host age revealed differences in neutrophil phenotype that suggest neutrophils from young mice or mild infections may have impaired bactericidal capacity and immune-suppressive functions compared neutrophil subsets from old mice or severe infections. Among CN, mild- and old-associated clusters P7 and P14 have less CD62L and more CD101 relative to the largest early severe SPN-associated CN cluster P3 (**Extended Data Figure 4AC**). CN (P14) and BALN (P19) from aged mice have lower expression of CD177 and Ly6G, but higher CD101 compared to comparable early and severe clusters (**Extended Data Figure 4A-C**). In BALNs, P1 is both an early and a mild associated cluster with high activation markers CD11b and CD177, as well as high PD-L1 and CD101 (**Figure 3CD, and Extended Figure 4C**). PD-L1 expression has recently been shown to be present on immune suppressive neutrophils and results in susceptibility to *P. aeruginosa* infection due to impaired phagocytosis capacity [29]. PD-L1 expression on neutrophils also promotes prolonged neutrophil survival and increases acute lung damage [30], and inhibit T cell function via ROS-mediated suppression and by suppressing T cell proliferation [31, 32]. This suggests that neutrophils from mild infections may be more immune suppressive than their counterparts from severe infections. Cluster P19 is enriched in BALN from aged mice and has high CXCR2 and CD101 expression relative to early (P1) and severe (P2) clusters but low PD-L1, CD18, CD177, and Ly6G (**Extended Figure 4BC**). This suggests that neutrophils from aged mice have this unique subset that are recently migrated and pro-phagocytosis. Taken together, these data suggests that mild infections result in active but more immune-suppressive neutrophils compared with severe infections, while neutrophils from old hosts tend to be less activated but more likely to be pro-phagocytosis compared to young.

**Figure 4.**
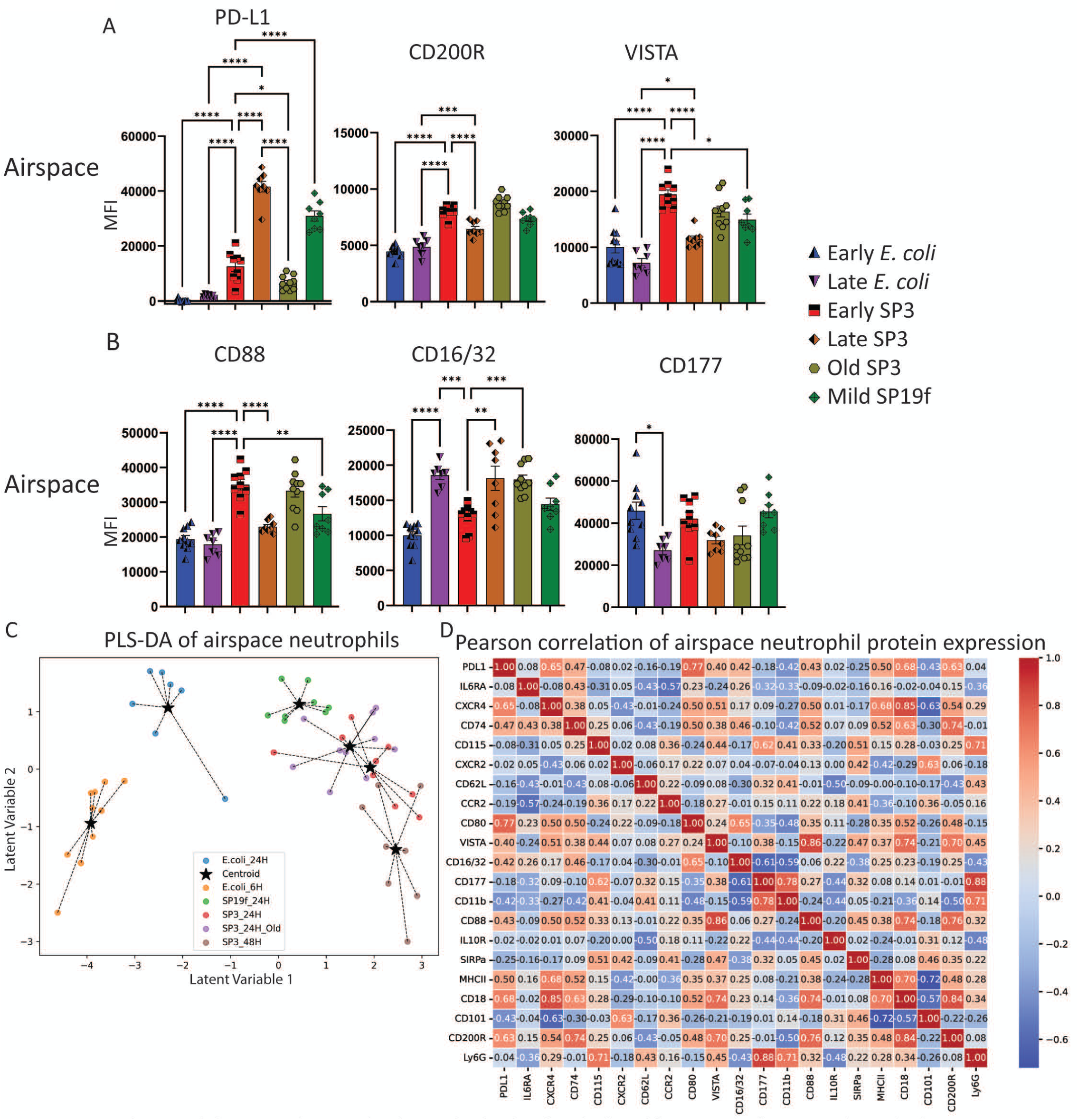
Immune regulators were highest on BALN from SPN-infected mice and track with markers of pathogen killing. Bar graphs of BALN groups show median fluorescence intensity (MFI) of (**A**) immune regulators or (**B**) markers of pathogen killing. Partial least squares discriminant analysis (PLS-DA) was used to classify all BALN samples based on average marker expression per subject and the (**C**) top two latent variables (components 1 and 2) are plotted. (**D**) PLS-DA loading values showing the contribution of each feature to the separation on each latent variable is also shown. (**E**) Pearson correlations (PC) or pairwise comparisons show strong (PC ≥ 0.7) and mild (PC ≥ 0.3) correlations between features. Strong positive correlations are indicated in red and strong negative correlations are in blue. Statistics used were one-way ANOVAs with Bonferroni multiple comparisons corrections where * = P < 0.05, ** = P < 0.01, *** = P < 0.001, and **** = P < 0.0001. Error bars show standard error of the mean (SEM).

### BALN phenotype is predictive of infection and time-point, and immune regulators CD200R and VISTA on BALNs correlate with markers of neutrophil degranulation

Next, we wanted to zoom in on differences between BAL neutrophil phenotypes, including some of the more subtle differences between aged and mild SPN infections. To investigate BAL neutrophil phenotype on a subset of proteins analyzed, we plotted the MFI of immune regulators (**Figure 4A**) and pathogen killing related proteins (**Figure 4B**) for all neutrophils from each experimental condition. Interestingly, PD-L1 was highest on late SP3 BALNs while CD200R and VISTA were highest on early SP3 infections (**Figure 4A**). VISTA is notable, as expression on neutrophils has been associated with diminished pro-inflammatory cytokine production in Lupus [33] but increased CD88 (C5aR) in Rheumatoid arthritis [34] and increased degranulation in crescent glomerulonephritis [35, 36]. Very recently, VISTA has been reported on neutrophils from patients with critically severe COVID-19 [37], which may suggest that it plays a regulatory role during pneumonia. Similarly, CD200R has been reported to affect neutrophil phagocytosis and ROS production [38], and exert other immune modulatory effects including enhancing Treg activity by CD200R^High^ neutrophils [39]. The role of immune regulators such as VISTA and CD200R on neutrophils will be further investigated in the next section.

To determine if total airspace (BAL) neutrophil phenotype stratifies by infection and time, partial least squares discriminant analysis (PLS-DA) was performed similar to previous work [40]. The top two latent variables (LVs) for BALN from each subject were plotted, demonstrating separation of the *E. coli* and SPN-infected groups (**Figure 4C**). Loading values for component 1, 2 and 3 are shown in **Supplemental Figure 2A**, demonstrating the specific features that distinguish the experimental groups. LV 1 separates infection and time post-infection groups well (**Figure 4C**), where CD101, CD11b, and CXCR2 have a negative loading score that distinguish *E. coli*-BALN and CD16/32, CD18, CD200R, CD74, CD80, CD88, CXCR4, IA/IE (MHCII), PD-L1, and VISTA stratify SPN-BALN (**Figure 4C; Supplemental Figure 2A**). Components two and three contain lesser degrees of variance (Loading values for each marker expression shows contribution to each latent variable: **Supplemental Figure 2A**), and the model accurately predicts which group each sample belongs to 80.77% of the time (cross-validation accuracy kfold = 80.77%; confusion matrix in **Supplemental Figure 2B**; and variable importance in projection score shows key model features in **Supplemental Figure 2C**).

Next, we were interested in identifying covarying protein expression to gain insights into possible mechanisms of neutrophil regulation. We thus used Pearson correlations (PC) of pairwise comparisons to determine features that covary and thus may be coregulated and have functional implications (**Figure 4D**). Ly6G,CD177, and CD11b have the strong correlations (highest correlation in the dataset is between Ly6G and CD177 PC = 0.88), though they appear to be expressed highest on activated early BALN phenotypes in *E. coli* and SPN-infected mice, possibly associated with activated and freshly demarginated neutrophils (**Figure 1G**; **Figure 4D**). CD80, CD74, CXCR4 and MHCII (only strong negative correlation) were positively correlated (PC ≥ 0.7), though their changes in expression between groups was modest at best, suggesting a strong mathematical but limited biological relevance (**Figure 4D**; **Supplemental Figure 2D**). Interestingly, immune regulators VISTA and CD200R strongly correlate (PC ≥ 0.7) with markers of CD88, CD18 and each other (VISTA and CD200R). C5a:C5aR stimulation can result in neutrophil degranulation and NETosis [41, 42], and CD16/32 activation can lead to degranulation and phagocytosis [43]. Taken together, the relationship between immune regulators VISTA and CD200R and neutrophil function worth investigating since increased expression of VISTA and CD200R correlates with markers of neutrophil bacteria killing (CD88 and CD18).

### Both VISTA and CD200R stimulation impede phagocytosis and enhance respiratory burst

Since less is known about the specific role of VISTA and CD200R on neutrophils, we sought to determine the functional impact of these receptors on neutrophil function via an *ex vivo* model system to testing phagocytosis, ROS production, and pathogen killing (schematic: **Figure 5A**).

**Figure 5.**
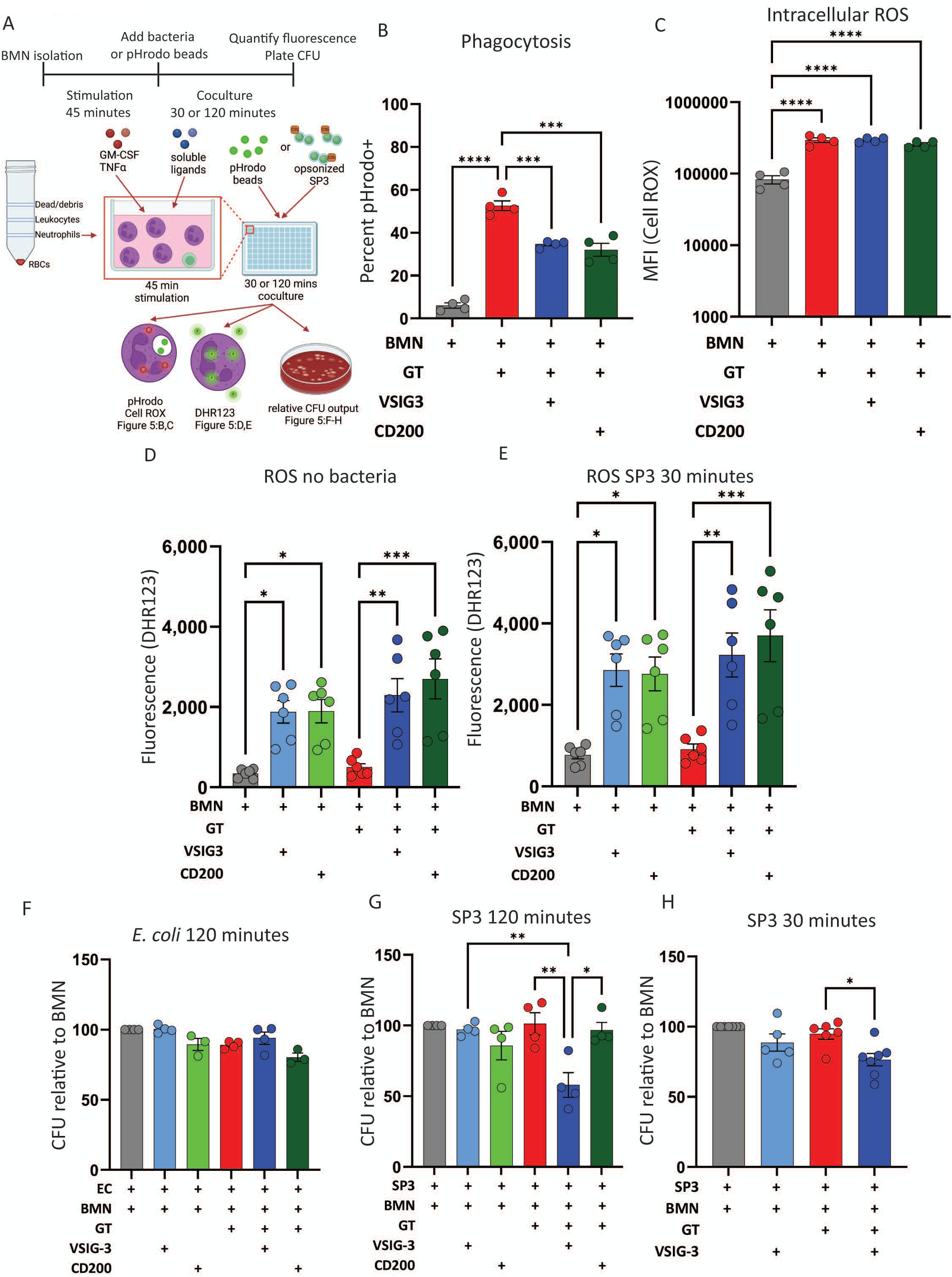
VISTA and CD200R stimulation of BMNs impedes phagocytosis and enhances ROS production, while VISTA stimulation enhances neutrophil killing of SP3. (**A**) Schematic representation of in vitro experimental pipeline. Bone marrow neutrophils (BMN) were co-stimulated with or without GM-CSF and TNFα (GT) and/or soluble ligands (VSIG-3 or CD200) for 45 minutes. Stimulated BMNs were then co-cultured with pHrodo beads or opsonized bacteria. Phagocytosis and intracellular ROS were analyzed via flow cytometry, intra-versus extra-cellular ROS was investigated by plate reader and CFU on blood agar plates. (**B**) Percent of pHrodo+ BMN or (**C**) intracellular ROS MFI across treatment conditions. DHR123 fluorescence after (**D**) 45 minutes ligand stimulation, or (**E**) co-cultured with SP3 for an additional 30 minutes. CFUs were counted and normalized to unstimulated BMN. Relative CFU after BMNs cocultured with (**F**) EC for 120 minutes, (**G**) SP3 for 120 minutes, or (**H**) SP3 for 30 minutes. Statistics used were one-way ANOVAs with Bonferroni multiple comparisons corrections or simple linear regression where * = P < 0.05, ** = P < 0.01, *** = P < 0.001, and ****= P < 0.0001. Error bars show standard error of the mean (SEM).

Bone marrow derived neutrophils (BMNs) were stimulated with GM-CSF and TNFα (GT, to activate neutrophils [44]) with or without soluble VISTA (VSIG-3) and CD200R (CD200) ligands [45–48]. We also initially investigated PD-L1 stimulation, but GT, fMLP, and immune regulator stimulation all failed to stimulate PD-L1 expression on BMNs (**Supplemental Figure 3A**), though VISTA and CD200R were both confirmed to be expressed on BMNs (**Supplemental Figure 3B**). Phagocytosis and intracellular ROS were analyzed via flow cytometry (gating strategy; **Supplemental Figure 3CD**) using pHrodo beads and Cell ROX, and total (intra- and extra-cellular) ROS production was analyzed using Dihydrorhodamine 123 (DHR123) via a plate reader (**Figure 5A**). GT stimulation significantly enhanced neutrophil phagocytosis of pHrodo beads, and co-stimulation with either VSIG-3 or CD200 impeded phagocytosis (**Figure 5B**). Conversely, VSIG-3 and CD200 treatment did not impact GT-induced production of intracellular ROS when analyzed by flow cytometry (**Figure 5C**). However, VSIG-3 and CD200 increase ROS above GT stimulation in the absence (**Figure 5D**) or presence of SP3 (**Figure 5E**). Taken together, these data suggests that CD200R and VISTA stimulation limit phagocytosis but enhance neutrophil extracellular ROS production independent of pathogen exposure.

### VISTA enhanced SP3, but not EC killing by bone marrow neutrophils

We next investigated the effect of VISTA and CD220R stimulation on neutrophil pathogen killing. Stimulated BMNs were co-cultured with opsonized bacteria for 30 or 120 minutes, and colony-forming units (CFU) were counted to determine bacteria killing (**Figure 5A**).

Interestingly, neutrophil stimulation with GT, VSIG-3, or CD200 alone had no impact on bacteria killing at any timepoint (**Figure 5FGH**). However, the combination of GT and VSIG-3 resulted in a significant increase in SP3 but not ED killing, while GT-CD200 did not affect killing (**Figure 5FGH**). Extracellular DNA, likely produced by BMN NETosis or lytic cell death, was measured using a non-cell-permeable DNA-binding dye (Cytox Green). The addition of the detergent saponin resulted in significant increases in Cytox Green fluorescence, but GT, VSIG-3, CD200, or the combination had no effect, both with and without bacteria (**Supplemental Figure 3F-G**). Taken together, these data suggests that VISTA stimulated neutrophils kill *S. pneumococcus* in a pathogen specific manner but were unlikely to do so via NETosis.

### VISTA stimulation results in altered surface marker expression and enhances degranulation

Given the positive correlation between VISTA and CD88, we further examined neutrophil degranulation in our model. BMNs were harvested and stimulated with GT, VSIG3 and/or CD200, supernatant was collected to perform ELISAs, and cells were stained and analyzed via flow cytometry. A modified version of the larger flow cytometry panel was designed to determine whether VISTA stimulation increases expression of markers associated with neutrophil degranulation (**Supplemental Table 3**). Neutrophils were again identified as CD45+ Ly6G+, live, single cells. First, we investigated the impact of VSIG-3 and CD200 stimulation on VISTA and CD200R expression. Heat inactivated (boiled) VSIG-3 was utilized as a negative control in all experiments and mirrors GT-BMN without VSIG-3 response (**Supplemental Figure 4A**). Curiously, GT and VSIG-3 co-stimulated BMNs had significantly higher levels of CD200R, while VISTA expression is unaffected by GT, VSIG-3, or CD200 stimulation (**Figure 6A**). Next, we looked at markers of degranulation with VISTA and CD200R stimulation. CD11b (multiple granules), CD177 (secondary granules), and CD63 (primary granules) all had significantly higher expression on GT-BMN with VSIG-3 co-stimulation compared to GT-BMN (**Figure 6BC**). We then decided to look at direct stimulation of degranulation. fMLP stimulation in neutrophils is reported to cause neutrophil degranulation [49] through multiple signaling pathways including PI3K and MAPK [50], though Vsir^-/-^ neutrophils still respond to fMLP [51]. We stimulated BMNs with fMLP with and without CD200 or VSIG3 and examined CD63 surface expression as a measure of degranulation. After 45 minutes, fMLP stimulation alone did not increase levels of CD63 BMNs, but co-stimulation of VSIG3 with fMLP led to higher levels of CD63 (**Figure 6D**). Additionally, GT-BMN cocultured with VSIG-3 had significantly higher levels of neutrophil elastase in the supernatant compared to GT-BMN alone, GT-BMN cocultured with inactive boiled-VSIG-3, or BMN alone (**Figure 6E**), further suggesting that VISTA stimulation results in release of neutrophil granule contents. Interestingly, VSIG-3 stimulation results in a significant decrease in CD101 expression (**Supplemental Figure 4AB**), which may reinforce our own results that suggest VISTA stimulation impedes phagocytosis (**Figure 5B**) and skews neutrophil pathogen killing towards degranulation and respiratory burst (**Figure 5DE**; **Figure 6B-E**). Taken together, these data suggests that VISTA stimulation on activated neutrophils results in degranulation independent of specific GM-CSF, TNFα, or fMLP stimulation alone.

**Figure 6.**
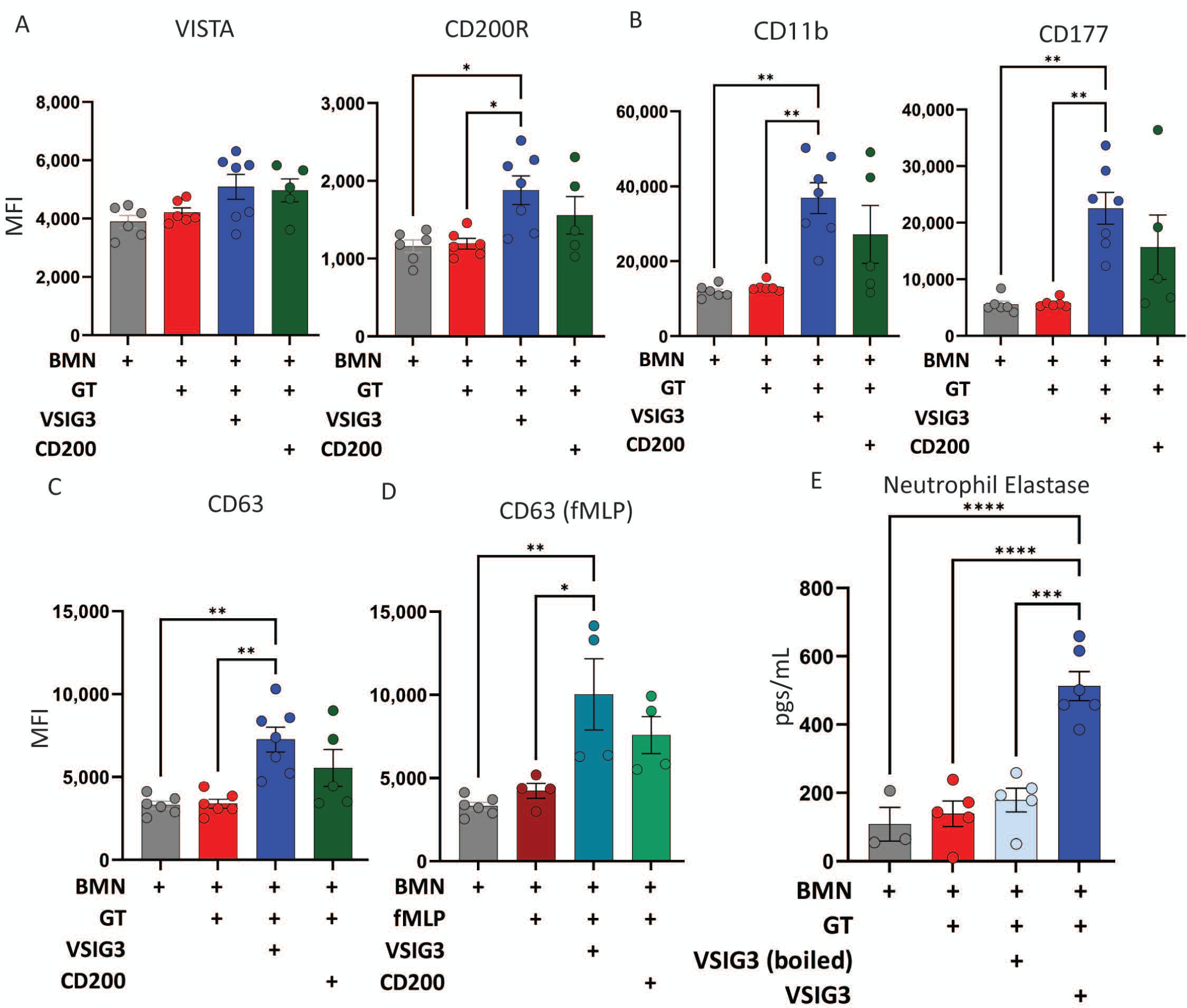
Co-stimulation of BMN with GT and VSIG-3 causes neutrophil degranulation. BMNs were stimulated with or without GT, fMLP, VSIG-3, heat inactivated (boiled) VSIG-3, or CD200. Bar graphs show flow cytometry data for GT-BMN of (**A**) immune regulators VISTA and CD200R, (**B**) markers of activation and degranulation CD11b and CD177, (**C**) degranulation CD63, (**D**) CD63 from fMLP-BMN. (**E**) The supernatant from GT-BMN was collected and neutrophil elastase was quantified via ELISA. Statistics used were one-way ANOVAs with Bonferroni multiple comparisons corrections or simple linear regression where * = P < 0.05, ** = P < 0.01, *** = P < 0.001, and **** = P < 0.0001. Error bars show standard error of the mean (SEM).

## Discussion

In this work, we comprehensively examined changes in neutrophil cell surface phenotype in response to several models of bacterial pneumonia. Our work uniquely identifies discrete circulating neutrophil surface protein phenotypes in response to distinct pathogens (*E. coli*, *S. pneumoniae*, *S. aureus*, and *P. aeruginosa*) during murine pneumonia with or without direct pathogen interactions. Altered activation state and phenotype in response to age, virulence, time post-infection, infection, and tissue were also observed. We also demonstrate that activation of the surface receptor VISTA, which we show is highest on BALN from SP3 infected mice, results in enhanced bacterial killing of SP3 but not EC, possibly secondary to enhanced degranulation but impaired phagocytosis. Our work suggests that VISTA stimulation results in neutrophils skewing away from phagocytosis towards degranulation, and that SPN infected mice have neutrophils consistent with a degranulation phenotype while EC infected mice have neutrophils with a more pro-phagocytosis phenotype. To our knowledge we were the first to report VISTA on neutrophils as a regulator that can cause pathogen specific degranulation and killing.

Both EC and SPN have a variety of immune escape mechanisms which impact neutrophil effector function. Our work is the first high-parameter (20+ color) flow cytometry study to directly compare SPN- and EC-directed neutrophil phenotypes and shed light on how these different immune escape strategies could result in differences in neutrophils based on different infections. SPN is known to evade phagocytosis [52] and induce high degrees of oxidative burst [53]. This may explain why SPN-BALN are PD-L1^High^, CD101^-^, and CD88^High^. PD-L1 has been shown to be on immune-suppressive neutrophils with impaired phagocytosis [29], CD101 has been shown to be on pro-phagocytosis neutrophils [20, 21], and CD88 (C5aR) is required for complement mediated activation and degranulation [27, 28]. On the EC side, hydrogen peroxide (H_2_O_2_; ROS species) has been shown to positively regulate neutrophil phagocytosis of EC through ERK and JNK pathway activation [54]. There is also some evidence that gram-negative bacteria (*P. aeruginosa* and *E. coli*) take advantage of phagocytosis to survive in the intracellular space and evade pathogen clearance, [55, 56] and tuning neutrophil function toward phagocytosis may be useful for EC survival, but dispensable for SPN. Further, phagocytosis of EC occurs in a CD18-dependent manner (results in CD18 internalization) [57], which may also explain why CD18 is drastically higher (and CD101 is drastically lower) on SPN-BALN compared to EC-BALN in our dataset. Taken together, our phenotyping data suggests that neutrophils in EC pneumonia are pro-phagocytosis (CD101^+^ CD18^Low^ PD-L1^-^) but pro- degranulation (CD88^High^ VISTA^High^ PD-L1^+^ CD101^-^) in SPN pneumonia.

VISTA (V-domain immunoglobulin suppressor of T cell activation, also known as PD-1H) is a B7 family member that is commonly expressed on lymphocytes, including macrophages and neutrophils [58]. VISTA can act as both a ligand and a receptor, and predominantly binds to either VSIG-3 (IGSF11) and PSGL-1 (at pH ∼6), with other less well confirmed ligands including Galectin-9, VSIG-8, metalloproteinase-13, syndecan-2, and leucine-rich repeats and immunoglobulin-like domains 1 (LRIG1) [48, 59, 60]. VISTA function on neutrophils is poorly understood, and has only recently been characterized in cancer [61] and autoimmune diseases [35]. Murine studies have shown that neutrophil and macrophage VISTA binding and activation during LPS challenge results in significantly lower inflammatory cytokine (IL-6, TNFα, and IL-12p40) production compared to LPS only stimulation [62] [33]. VISTA KO (*Vsir*^-/-^) mice have augmented TLR-mediated proinflammatory cytokine production compared to WT mice [63], suggesting an inhibitory role for VISTA on myeloid cells. Paradoxically, in human inflammatory diseases, neutrophil expression of VISTA is associated with lower chemokine and inflammatory cytokine production in Lupus but may promote inflammation via increased degranulation in crescentic glomerulonephritis, and increased C5aR expression in Rheumatoid arthritis [35]. In other whole-body VISTA KO (*Vsir*^-/-^) murine autoimmune models, *Vsir*^-/-^ myeloid cells have been shown to have increased migration (granulocytes and monocytes) [33], lower expression of C5aR (macrophages) [34], and lower degranulation (neutrophils) via inhibition of immune-complex activation [51]. Although some recent work has characterized how whole-body VISTA deficiency (*Vsir^-/^*^-^) impacts neutrophil function in autoimmune diseases, VISTA function on neutrophils during infection is poorly understood. To our knowledge, this is the first work that characterizes VISTA expression during bacterial pneumonia and shows differences in pathogen-specific killing of bacteria by neutrophils.

Despite commonly being referred to as an “inhibitory receptor,” our work shows that VISTA stimulation results in a protective activating role on neutrophils during pneumonia. Here, we show that VISTA expression is higher on BALN from mice infected with discrete pathogens *in vivo*, and that VISTA stimulation results in pathogen-specific killing and degranulation *in vitro*. Very recent work on severe COVID-19 patients also associates high VISTA expression with severe disease and C5aR1 and CD177 expression, which agrees with our own data and suggests that our own finding may be applicable to human biology and disease as well [37]. Interestingly, VISTA stimulation also results in significant phenotypic changes, including higher markers of degranulation (CD11b, CD66a, CD177, and CD63), CD200R, as well as and lower CXCR2.

Enhanced CD200R expression upon VISTA stimulation is notable as CD200R may also be a viable therapeutic target, and CD200R stimulation results in enhanced respiratory burst as seen in our *in vitro* data. CD101 expressing neutrophils were better at phagocytosing and clearing *Salmonella enterica* compared to Cd101^-/-^ neutrophils [19], which may further support our argument that VISTA stimulation skews neutrophils from phagocytosis towards degranulation and respiratory burst.

Both recent reports and our own work indicate that VISTA may be a therapeutic target during pneumonia. VISTA stimulation enhances neutrophil degranulation while simultaneously enhancing expression of CD200R, both of which enhance respiratory burst and could lead to faster clearance of bacteria, particularly in SPN infections. While degranulation and respiratory burst have the potential to contribute to host tissue damage, VISTA stimulation is also known to reduce inflammatory cytokine production [33, 62], and thus may also reduce tissue damage.

VISTA targeted therapeutics are in development, particularly as a blocking anti-VISTA (or anti-VSIG-3 or -PSGL-1) which works synergistically with anti-PD-L1, anti-PD-1, or anti-CTLA-4 treatment in various cancers which result in increased inflammation and improved outcomes [60, 64–66]. To our knowledge, no activating VISTA therapeutic has been attempted, though further work is needed to determine the viability of VISTA activating therapies during bacterial pneumonias.

CD200R is expressed ubiquitously on neutrophils and macrophages, and binding of soluble or cell-bound CD200 results in Erk signaling [67–69]. In *in vitro* studies, human macrophages stimulated with CD200 have less phagocytic capacity and inflammatory cytokine production (TNFα) compared to untreated [70]. In whole body CD200R KO studies of *F. tularensis* lung infections, CD200R KO mice have neutrophils with impaired ROS production and higher bacterial burden compared to wildtype [38]. Similarly in a *B. pseudomallei* lung infection model, CD200-Fc treated mice have enhanced survival compared to untreated [71]. Here, we show enhancement of CD200R expression on BALN in a pathogen specific manner, and that CD200R stimulation in vitro impedes BMN phagocytosis and enhances respiratory burst. Taken together, our work adds to the current body of literature that suggests that CD200R may be ideal as a therapeutic target in acute lung infections.

Neutrophil dysfunction associated with advanced age has been investigated by numerous groups. Notably, neutrophils from older subjects were known to have dysfunctional recruitment, resolution of inflammation, and effector function, including impaired microbe killing, phagocytosis, degranulation, and NETosis [72]. Phenotypic changes on neutrophils also include lower CD16 (but more CD16^high^ neutrophils), higher CD63 and CD11b, and lower CD73 expression [72]. In our data, aged mice have slightly lower levels of activated neutrophils (P3 and P2) which were relatively high for combinations of CD11b, CD177, Ly6G, and CD88 with corresponding higher levels of less activated neutrophils (P14 and P19) CN and BALN. Mild infection and advanced age also express modestly higher CD101 compared to neutrophils from young mice with severe pneumonia, which may suggest that these conditions favor phagocytosis over other degranulation and NETosis. Paradoxically, neutrophils from aged individuals are known to have less phagocytic capacity compared to those from younger individuals [73]. This may suggest that phagocytosis is more favored for clearing SPN in aged mice, though total killing capacity may be diminished. Neutrophil function in aged populations and the function of CD101 on neutrophils should be further investigated. Notably, neutrophils from aged mice also have modestly lower PD-L1 expression but comparable levels of CD200R and VISTA compared to young, which may suggest that these markers persist throughout life and may be viable therapeutic targets, which will be highlighted later in this discussion.

Overall, our work is a well-balanced characterization of circulating and BAL neutrophils during multiple types of bacterial pneumonia and identifies VISTA as a potential therapeutic target, though there were a few limitations that should be acknowledged. Functional studies were performed on BMNs due to the technical difficulty of isolating sufficient numbers of BALNs.

Further work should be performed to determine if VISTA stimulation on BALNs in vivo behave similarly to BMN *in vitro*. Soluble ligands were also used to assess VISTA and CD200R activation, which may miss cell-to-cell interactions that would be accompanied with other co-stimulatory binding, though there is evidence that soluble ligands for both exist in the lung and in peripheral blood [74–76].

Accumulating evidence suggests that neutrophils are highly dynamic immune cells that respond specifically to their environment. We demonstrate and add to this idea by showing that neutrophil phenotype is influenced by numerous variables including pathogen, tissue, time-point during infection, and age. We offer novel insights into specific phenotypes that would be clinically relevant to diagnosing specific types of bacterial pneumonia from circulating neutrophils. We also identify potential therapeutic targets, particularly VISTA. We showed that VISTA expression is highest on BALN from SP3 infected mice, and that VISTA stimulation enhances degranulation and killing of SP3 but not EC. Honing our understanding of neutrophil regulation during pneumonia in response to different pathogens is crucial, and VISTA should be further investigated as a potential therapeutic target during pneumonia.

## Methods

### Sex as a biological variable

Flow cytometry and in vitro studies were performed on roughly equally numbers of male and female mice. Sex was investigated as a variable, though no significant differences in neutrophil phenotype nor function were identified.

### Murine model of pneumonia and tissue collection

C57BL/6J mice were purchased from Jackson Laboratories (Jax.org). Mice were between 6-12 weeks of age, and aged mice were 90-100 weeks old and were aged in-house. Mice were housed in a barrier facility and provided food and water *ad libitum*. For pneumonia models frozen bacteria was cultured on blood agar plates for 10-12 hours *Streptococcus pneumonia* (SPN; 1.7-2.8*10^6^ CFU SPN, serotype 3 (SP3), strain catalog number 6303 ATCC; or 0.2-1.7*10^6^ CFU serotype 19f (SP19), strain EF3030 [77]) or 1.6-2.4*10^6^ CFU *Escherichia coli* (EC; serotype 06:K2:H1; American Type Culture Collection [ATCC] no. 19138) was intratracheally injected into the left lung lobe of mice as previously reported [78]. For low dose infections, EC, SP3, *Staphylococcus aureus* (strain 25923 ATCC) and *Pseudomonas aeruginosa* (PA01; gifted to Dr. Joseph Mizgerd by Dr. Alice Prince) were cultured on blood agar plates from frozen aliquots and allowed to grow to stationary phase (16 hours (24 hours for PA01) and 5 to 20*10^4^ CFU (or 1-2*10^6^ CFU for *S. aureus*) was intratracheally injected into the left lung lobe of mice for 24-hours.

Mice were euthanized and tissue was collected 6-, 24-, or 48-hours post infection. Blood was collected from the inferior vena cava using a heparinized needle. An aliquot of whole blood was serially diluted and plated on blood agar to determine blood CFU. The remaining blood had red blood cells removed using RBC Lysis Buffer (Biolegend), was washed in PBS, and kept on ice until ready for staining. For bronchioloalveolar lavage (BAL), the heart-lung block was surgically harvested and suspended in air by the trachea attached to a blunt catheter, and serially lavaged with 1 mL ice-cold PBS, for a total volume of 10 mL. The first mL BALF had cells removed by centrifugation (350 relative centrifugal force (RCF), 10 minutes) and the supernatant was frozen at -80°C for total protein other quantitative analyses. The remaining BALF (approx. 9ml) was centrifuged (350 RCF, 10 in), cells pooled together, resuspended in PBS and held on ice until stained.

### Spectral flow cytometry

Similar to our previous work [79], isolated neutrophils were washed and stained in phosphate buffer solution (PBS) with Live/Dead Blue for 15 minutes on ice in the dark, then washed with fluorescence activated cell sorting (FACS) buffer (PBS with 2% fetal bovine serum (FBS) and 2 mM ethylenediaminetetraacetic acid). Cells were kept on ice and in the dark throughout, and in low-stick polypropylene V-bottom (Corning; non-tissue culture treated) plates for staining and in polypropylene cluster tubes (Corning) for acquisition. Cells were first stained with CD16/32-Brillinant Violet 786 for five minutes, then cells were blocked with mouse Fc-blocking reagent (Biolegend) and stained with the additional 22-color antibody reagent cocktail. The antibody cocktail (**Supplemental Table 1**) and fresh single stain controls were prepared fresh for each experiment by pipetting in-house titrated antibodies to FACS buffer with BD Brilliant Stain Buffer (BD Biosciences). Cells were stained in 100 µLs, incubated for 20 minutes, washed twice with FACS buffer, and kept on ice and in the dark until acquired on a 5-laser Cytek Aurora (Cytek Biosciences). Each experimental group was split across at least two different experimental runs over eight total experimental runs.

### Analysis pipeline for flow cytometry

Daily QC was run on the 5-laser Aurora (Cytek Biosciences) which automatically adjust the gains (voltages) of each of the 64 raw channels to ensure consistency of median fluorescence intensity (MFI) across experimental days and control for batch. Samples were recorded using SpectroFlo version 2.2 (CytekBioscience) and unmixed using the integrated ordinary least square spectral unmixing algorithm. Both expert-guided and algorithmic data analyses were performed using Omiq.ai cloud analysis platform.

Live, single cell, CD45+ Ly6G+ cells (neutrophils) were identified via manual expert-guided gating and further unbiased analysis using FlowAI [80], opt-SNE [81], and PhenoGraph [82] (gating strategy shown in **Supplemental Figure 1A**). Data were Arcsinh transformed (cofactor = 6000) and an equal number of cells (maximum equal per group) per experimental group were subsampled (maximum of 22,000 per sample) were concatenated for a total of 1,740,551 cells.

The concatenated and subsampled file was used to produce an opt-SNE map with PCA-pre-initialization, and the PhenoGraph clustering algorithm (kNN = 20, distance metric = Euclidean) was used as an unsupervised method to automatically subset our data using a curated subset of features that reduces noise in clustering. For both opt-SNE and PhenoGraph, the clustering parameters used were SSC-B, PD-L1, CD74, CXCR2, CD62L, CCR2, CD80, VISTA, CD16/32, CD177, CD11b, CD88, SIRPα, IA-IE (MHCII), CD18, CD101, CD200R, and Ly6G. 37 Distinct PhenoGraph clusters were identified and two small clusters that were >50% from a single subject were excluded from these analyses. The 35 clusters were overlayed in unique colors on an opt-SNE map (**Figure 1B**). MFI of samples, MFI of clusters, and per cent of total neutrophils contributing to each PhenoGraph cluster were exported from Omiq.ai and analyzed in Prism 10.0 (Graphpad).

To ensure that run-to-run variation was minimized, uninfected circulating neutrophils were stained with the full panel and each channel was compared using offset histograms for each channel on each experiment run (example channels, **Supplemental Figure 1B**). Each experiment run within each experimental group was also compared using offset histograms for each channel to further monitor run-to-run variation (example of five BAL groups, **Supplemental Figure 1C-G**). Across both strategies for monitoring batch effects, some heterogeneity within batches was observed, though differences in MFI between experimental runs was negligible.

### Bone marrow neutrophil isolations

Bone marrow neutrophil (BMN) isolation was adapted from Ubags and Suratt (2018) [83]. In brief, femurs and tibias were removed from euthanized mice and flushed with HBSS with 0.38% Sodium Citrate (HBSS-SC) until the bone was blanched. BM was filtered with a 40-micron filter and washed with HBSS-SC. Percoll (Fisher Scientific) density gradient layers of 72%, 64%, and 52% (‘100%’ Percoll (90% Percoll, 10% 10x HBSS)) were diluted with HBSS-SC, and the washed and filtered BM was layered on top and spun at 1550g for 30 minutes with no brake.

Debris and lymphocyte layers were removed first to minimize contamination, then the neutrophil layer was extracted, washed, and analyzed for cell count, viability, and Cytospin with differential cell counts was performed to confirm the fraction of granulocytes that were isolated (must be >80% granulocytes to be used for experiments, typical purity was 85-95%). For BMN stimulation,5*10^5^ BMN in 50 µL of DPBS++++ per well were stimulated for 45 minutes at 37°C in the dark with TNFα (R&D; *E.coli*-derived; 20 ngs/mL), GM-CSF (R&D; *E. coli*-derived; 50 ngs/mL), VSIG-3 (R&D; mouse myeloma cell line (NS0)-derived; 10 µgs/mL), CD200 (Biolegend; 293E-derived; 10 µgs/mL), and/or PD-1 (R&D; mouse myeloma cell line (NS0)-derived; 10 µgs/mL).

### Bacterial clearance and extracellular DNA assays

Bacterial clearance assays were performed using BMN and an adapted protocol from by Dr. Heidi Welch [44]. In brief, BMN were isolated as discussed in the previous section, resuspended in DPBS++++ (Ca^2+^, Mg^2+^, 0.1% glucose, 4 mM NaHCO_3_), and were stimulated with or without GM-CSF + TNFα (GT) and soluble ligands against immune regulators for 45 minutes at 37°C in the dark in 50 µLs in white-walled clear flat-bottom 96-well plates (Corning). Bacteria (same SP3 or EC used in *in vivo* infections) were diluted in ice-cold DPBS++ (Ca^2+^, Mg^2+^) and kept on ice until they were co-cultured with the stimulated BMNs at an MOI of 1 (0.5 – 1.5) at 37°C in the dark for 30 or 120 minutes. Cultures were mixed with a pipette and plated on blood agar to count CFUs. For the 120 minute clearance assays, CYTOX Green (Thermo Fisher Scientific) was added to the cocultured bacteria and BMN to measure extracellular DNA, and fluorescence was read on a Synergy LX plate reader (BioTek) prior to CFU plating.

### Phagocytosis and ROS productions assays

BMN isolation and stimulations were performed similarly to the previous section. For phagocytosis and intracellular ROS assays, a cocktail of pHrodo Red *E. coli* BioParticles (Thermo Fisher Scientific) and CellROX Green (Thermo Fisher Scientific) were added to stimulated BMNs for 30 minutes at 37°C in the dark, washed twice, and resuspended in FACS buffer with 7-AAD. Cells were kept on ice in the dark until they were acquired in the same 5-laser Aurora (Cytek Biosciences) as previously discussed. BMN were gated using FSC vs SSC, live (7-AAD negative) single cells, and pHrodo positive gate was set using a fluorescence minus one (FMO) (**Supplemental Figure 2CD**). Anti-Ly6G-APC Fire 810 was stained to confirm Ly6G^+^ cell position on FSC and SSC (**Supplemental Figure 2E**).

For total ROS assays, BMN isolation and stimulations were performed similarly to the previous section, with the addition of Dihydrorhodamine 123 (DHR123) added to the BMN stimulation. Fluorescence was analyzed via a BioTek Synergy LX plate reader (Agilent) using Gen5 acquisition software before and after the 45 minutes stimulation, and after 30-minute coculture with SP3 for total ROS production.

#### ELISA

Mouse neutrophil elastase ELISA (R&D Systems) was performed per the included protocol. In brief, the capture antibody was incubated on the included plate overnight at room temperature. The next day, the plate was washed and blocked with the included reagent dilutant. Samples or standards were added to the appropriate wells and incubated for two hours before the detection antibody was added and incubated for an additional two hours with washes in between. Plates were washed again and Streptavidin-HRP was added to each well and incubated for 20 minutes, followed by washing, then substrate solution was added to each well for 20 minutes, followed by stop solution. The optical density (OD) of 450 nm and 540 nm were read via BioTek Synergy LX plate reader (Agilent) using Gen5 acquisition software and wavelength correction was performed by subtracting the 540 nm measurement from the 450 nm measurement for each well.

### Partial Least Squares Discriminant Analysis (PLS-DA) and Model Validation

All BALN samples were gated on live, CD45^+^, Ly6G^+^ single cells and MFIs for each subject were exported in a .csv file for further analysis. All data processing, modeling, and statistical analyses were conducted using Python (v3.13) using the following libraries: scikit-learn for PLS-DA, LASSO regression (data not shown), cross-validation, and classification metrics, numpy and pandas for data manipulation and numerical operations, matplotlib and seaborn for visualization of PLS scores, feature loadings, and model performance, and scipy.stats for statistical significance testing. Partial least squares discriminant analysis (PLS-DA) was performed to classify samples based on the measured features. Prior to modeling, all data were transformed using the inverse hyperbolic sine (arcsinh) function to stabilize variance across measurements and accommodate data skewness. Transformed data were then standardized by Z-score normalization across each feature. Cross-validation was performed using a Stratified K-Fold approach using 1/5 of the dataset to ensure balanced representation of each class within training and validation sets. The number of latent variables (LVs) was determined by iteratively testing models with up to 10 LVs and selecting the number that minimized mean squared error (MSE) across validation folds.

Model performance was evaluated by calculating classification accuracy from cross-validation, as well as visualizing separation in PLS score space using convex hulls and star plots.

Variable importance was assessed using PLS loadings and Variable Importance in Projection (VIP) scores. Features with high VIP scores (>1.0) were considered strong contributors to class separation. Additionally, Least Absolute Shrinkage and Selection Operator (LASSO; data not shown) regression was performed iteratively across all features to identify the most frequently selected predictors for classification. To assess the statistical significance of the classification model, permutation testing (n = 100) was performed by randomly shuffling class labels and rebuilding the PLS-DA model for each iteration. The distribution of errors from these randomized models was compared to the observed classification error using a Mann–Whitney U-test, with significance defined as p < 0.05.

### Statistics

Unless otherwise stated, data presented in figures were from at least two separate experiments. Unless otherwise stated, error bars were used to report standard error of the mean in figures. The statistical method used to compare multiple treatment groups was a one-way ANOVA with Bonferroni multiple comparisons corrections or simple linear regression. Asterisks for adjusted p values were * = P < 0.05, ** = P < 0.01, *** = P < 0.001, and **** = P < 0.0001. Statistical assumptions were made that data is normally distributed and that each observation in the dataset is independent of other observations.

## Declarations

### Ethics approval

All procedures involving animals were in compliance with BU IACUC

All animal studies presented in this manuscript were overseen by the Boston University Institutional Animal Care and Use Committee (IACUC). All protocols were approved prior to the performance of experiments (BU IACUC protocol #PROTO201800710).

### Data availability

Our study does not contain any datasets that are required to be uploaded to publicly available repositories, however our flow cytometry dataset may be made available to specific researchers upon request.

### Software for data analysis

For data analysis, commercial platforms such as OMIQ.ai and Graphpad Prism 10 were used for analyses. A custom code produced on Python by Dr. Anna Belkina was used to perform partial least squares discriminant analysis and model validation which is available on Github <https://github.com/ACBelkina/neutrophils_in_pneumonia>.

## Funding

RMFP: T32 HL 7035-48.

KET: R01HL158732, K08130582, L30 HL138777, F32 HL120551, T32 HL703547, KL2 TR001411.

Supported in part by a research grant from Drs. Margaret Seton and Joseph Jacobson in memory of Dr. Dennis J Beer.

### Author contributions

RMFP: Study design, experimental design, conducting experiments, and manuscript writing and preparation, KRM: conducting experiments, YL: experimental design and conduction experiments, LP: conduction experiments, LJQ: study design and experimental design, JPM: study design and experimental design, ACB: experimental design, and KET: study design, experimental design, and writing.

## Acknowledgements

We would like to thank Dr. Heidi Welch for sharing her bacterial clearance assay protocol with us and providing feedback and advice to make this assay work.

---

**Extended Data Figure 1.**
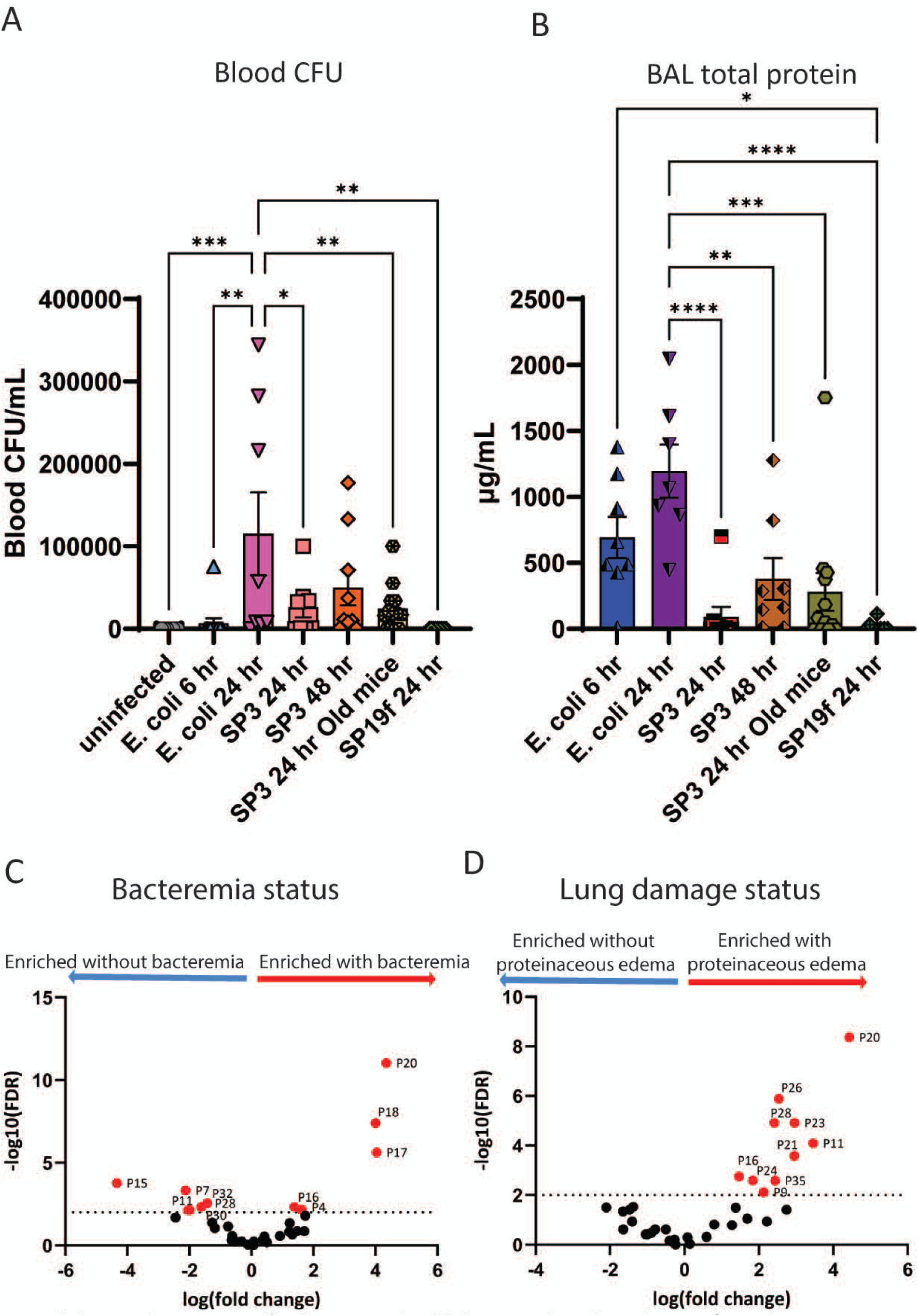
Specific clusters track with bacteremia or lung damage (proteinaceous edema) status. Bacteremia and total protein were quantified to assess physiological effects of bacterial pneumonia. (**A**) Blood CFUs from each mouse were calculated and plotted, along with (**B**) BAL total protein. To allow for bimodal comparisons, bacteremia status (0-1 colonies = no bacteremia, 2 or more = bacteremia) and lung damage observed via protein in the lung (protein detectable or not detectable) were defined. Clusters associated with mice that had (**C**) bacteremia or (**D**) measurable proteinaceous edema were plotted on volcano plots. Statistics in bar graphs used were one-way ANOVAs with Bonferroni multiple comparisons corrections or simple linear regression where * = P < 0.05, ** = P < 0.01, *** = P < 0.001, and **** = P < 0.0001. Error bars show standard error of the mean (SEM).

**Extended Data Figure 2.**
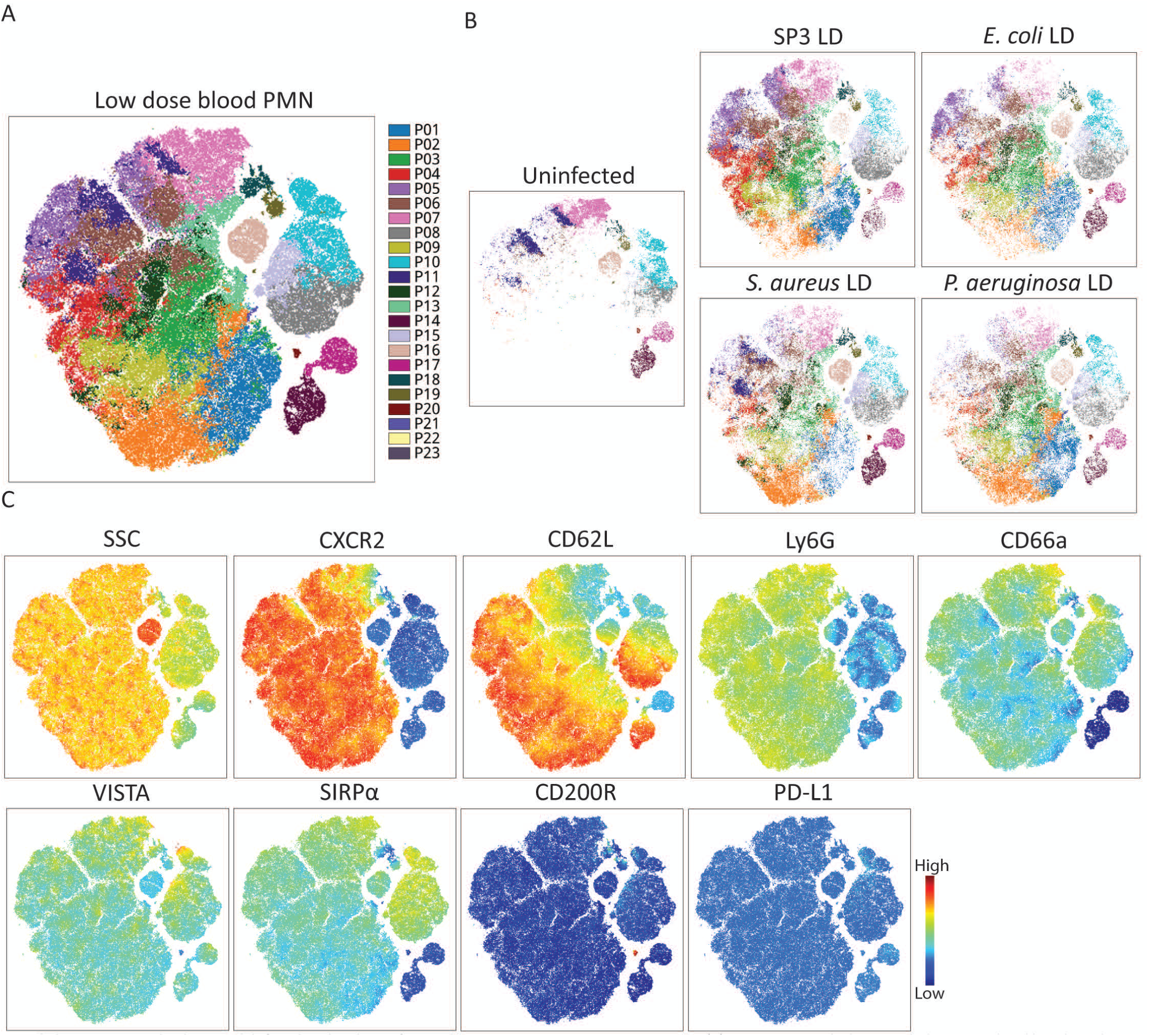
Blood neutrophils from low dose lung infections have unique protein expression patterns. (**A**) Opt-SNE map with PhenoGraph clusters overlayed by color. Subsets of the opt-SNE plot were also plotted to show where cells from (**B**) different low dose infections. (**C**) Heatmap overlays for examples of features that distinguish the different infection groups were plotted.

**Extended Data Figure 3.**
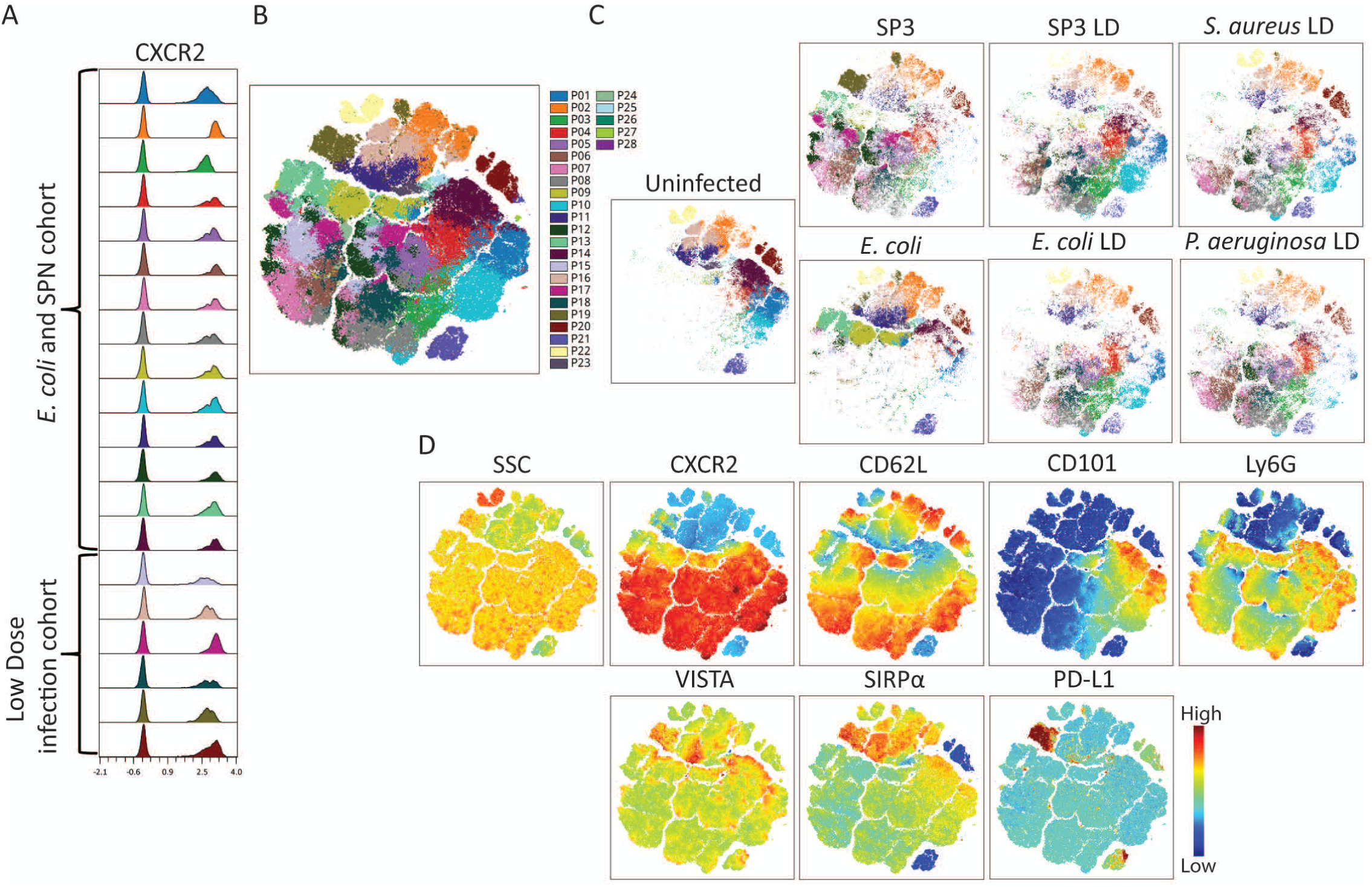
Blood neutrophils from lethal and low dose lung infections have unique protein expression patterns. To ensure compatibility between panels, CytomNorm was used to normalized channels, and an (**A**) example channel across all runs used is shown to demonstrate minimal variance. (**B**) Opt-SNE map with PhenoGraph clusters overlayed by color. Subsets of the opt-SNE plot were also plotted to show where cells from (**C**) different low dose infections. (**D**) Heatmap overlays for examples of features that distinguish the different infection groups were plotted.

**Extended Data Figure 4.**
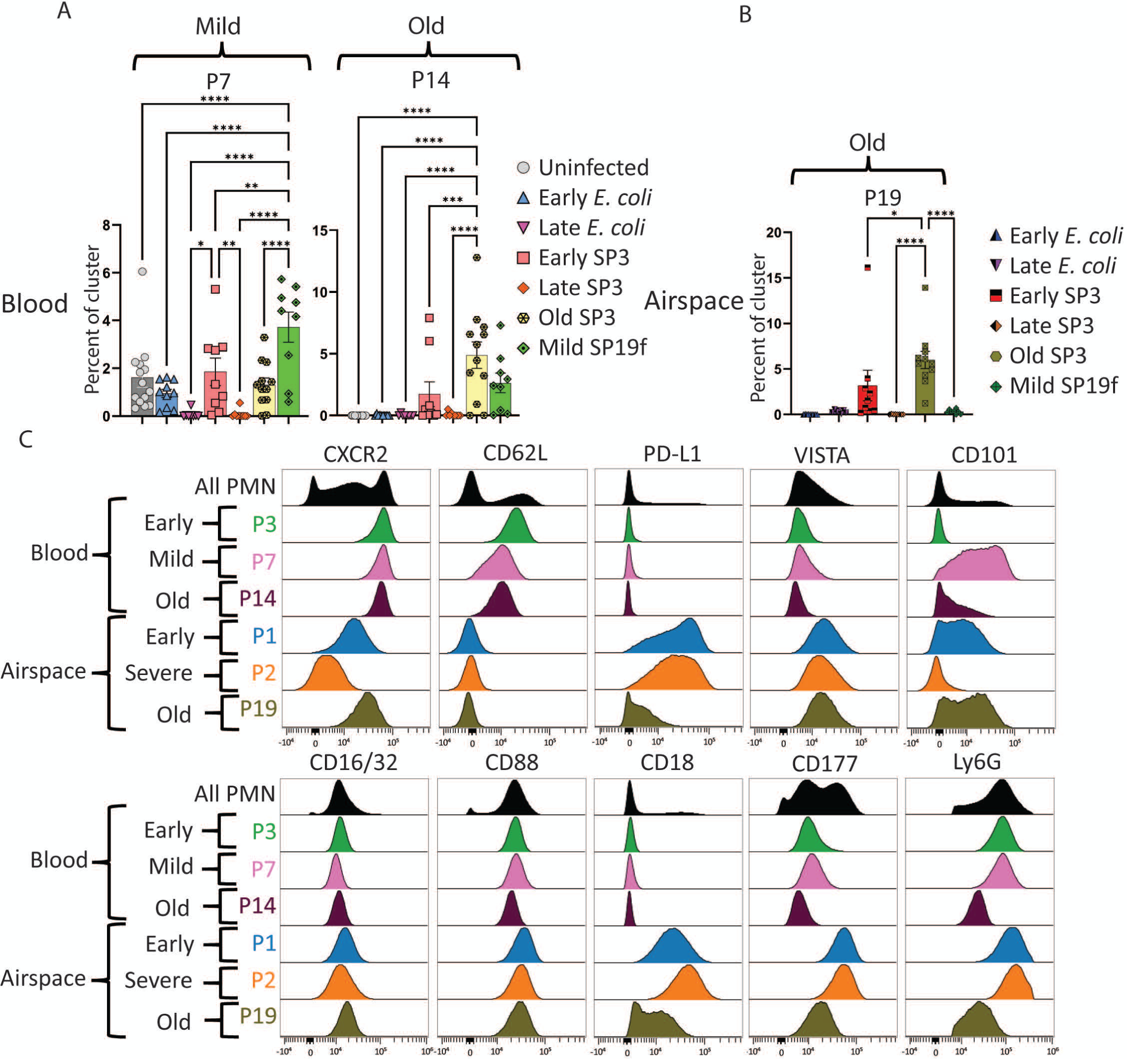
Impact of mild virulence and host age on neutrophil phenotype. Bar graphs show clusters with a significantly higher fraction of blood neutrophils from mice infected with (**A**) mild (P7) or from aged mice (P14), or from the BALF of (**B**) aged mice. Histograms show specific marker expression of each mild and age cluster relative to severe and young phenotypes respectively (**C**). Statistics used were one-way ANOVAs with Bonferroni multiple comparisons corrections where *= P < 0.05, ** = P < 0.01, *** = P < 0.001, and ****= P < 0.0001. Error bars show standard error of the mean (SEM).

**Supplemental Figure 1.**
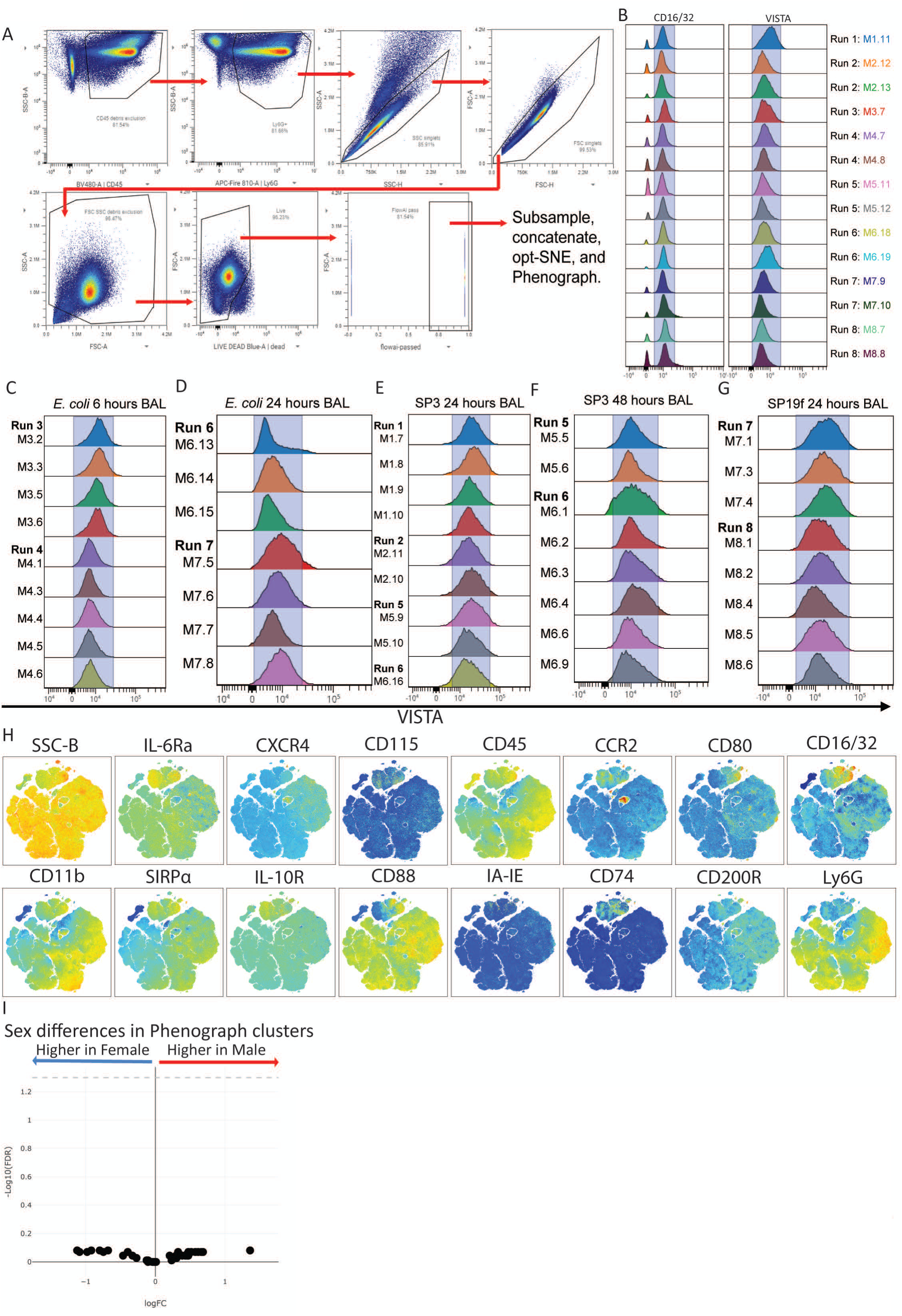
(**A**) Example of gating strategy for large spectral cytometry dataset. Cells gated were CD45+, Ly6G+, single cells, small debris excluded, live, that passed FlowAl analysis. There were eight total experimental days (runs; where MX.Y = Mouse_run#.mouse#), and as part of our batch effect quality control, uninfected blood was collected and stained for our panel on each run to compare changes in MFI. (**B**) CD16/32 and VISTA were shown as examples across all eight experimental days (runs) with a blue semi-transparent box along the positive to assist with visualization of consistency. All experimental groups were run on at least two experimental days, and VISTA expression for BALN experimental groups were shown with each sample across multiple runs for (**C**) EC 6-, (**D**) EC 24-, (**E**) SP3 24-, (**F**) SP3 48-, and (**G**) SP19f 24-hours. (**H**) Heatmap overlays of each parameter from Figure 1. (**I**) Sex differences were analyzed across all PhenoGraph clusters to determine if specific neutrophil phenotypes were enriched in male or female mice. Statistics used were one-way ANOVAs with Bonferroni multiple comparisons corrections where *= P < 0.05, ** = P < 0.01, *** = P < 0.001, and ****= P < 0.0001.

**Supplemental Figure 2.**
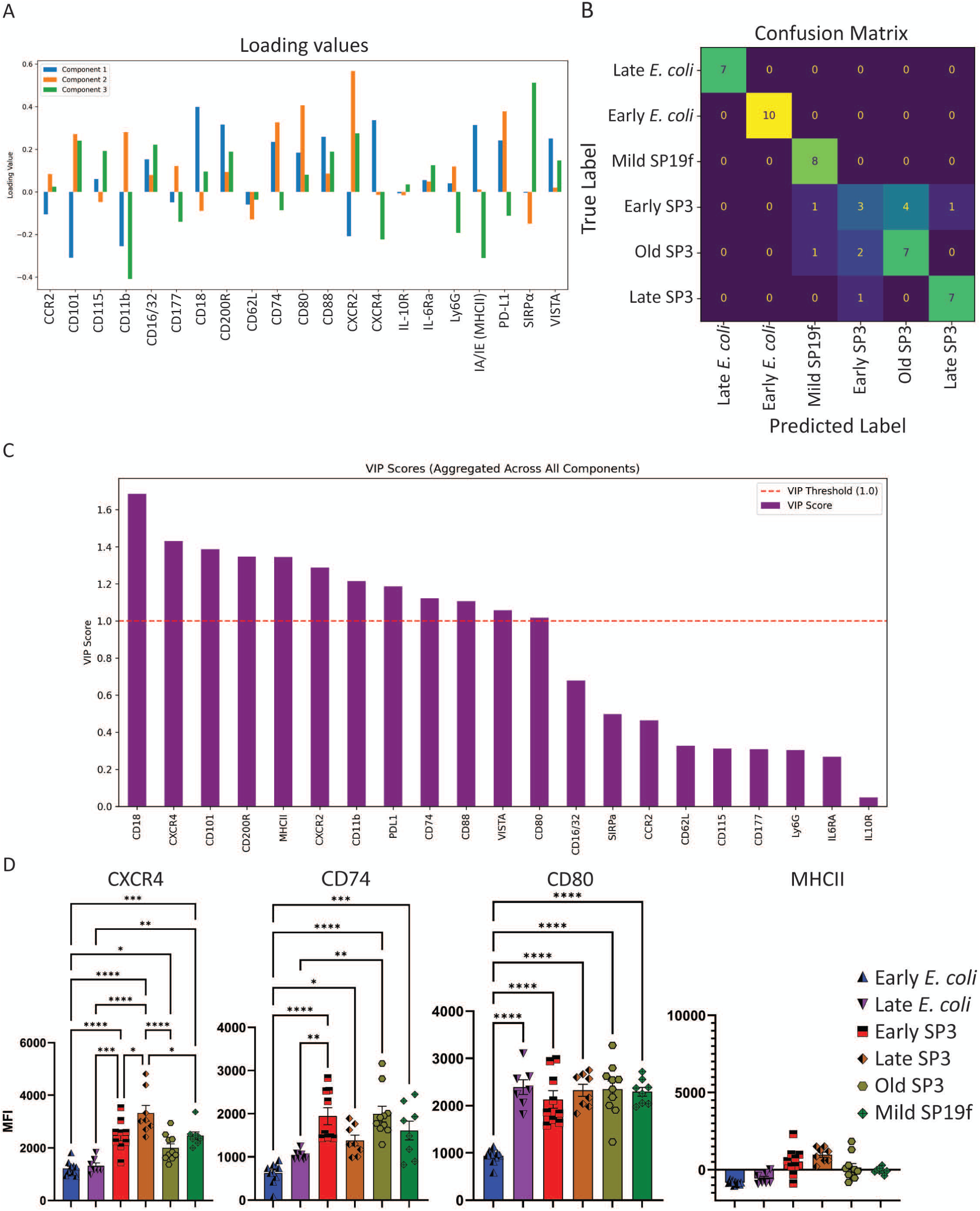
For PLS-DA analyses, (**A**) the PLS-DA loading values showing the contribution of each feature to the separation on each latent variable is also shown. (**B**) Confusion matrix and the (**C**) variable importance in projection (VIP) scores are shown where a VIP score greater than 1 significantly contributes to the model accuracy. (**D**) Bar graphs of BALN groups show median fluorescence intensity (MFI) for specific markers with strong Pearson correlation coefficients from Figure 4E. Statistics used were one-way ANOVAs with Bonferroni multiple comparisons corrections where *= P < 0.05, ** = P < 0.01, *** = P < 0.001, and ****= P < 0.0001. Error bars show standard error of the mean (SEM).

**Supplemental Figure 3.**
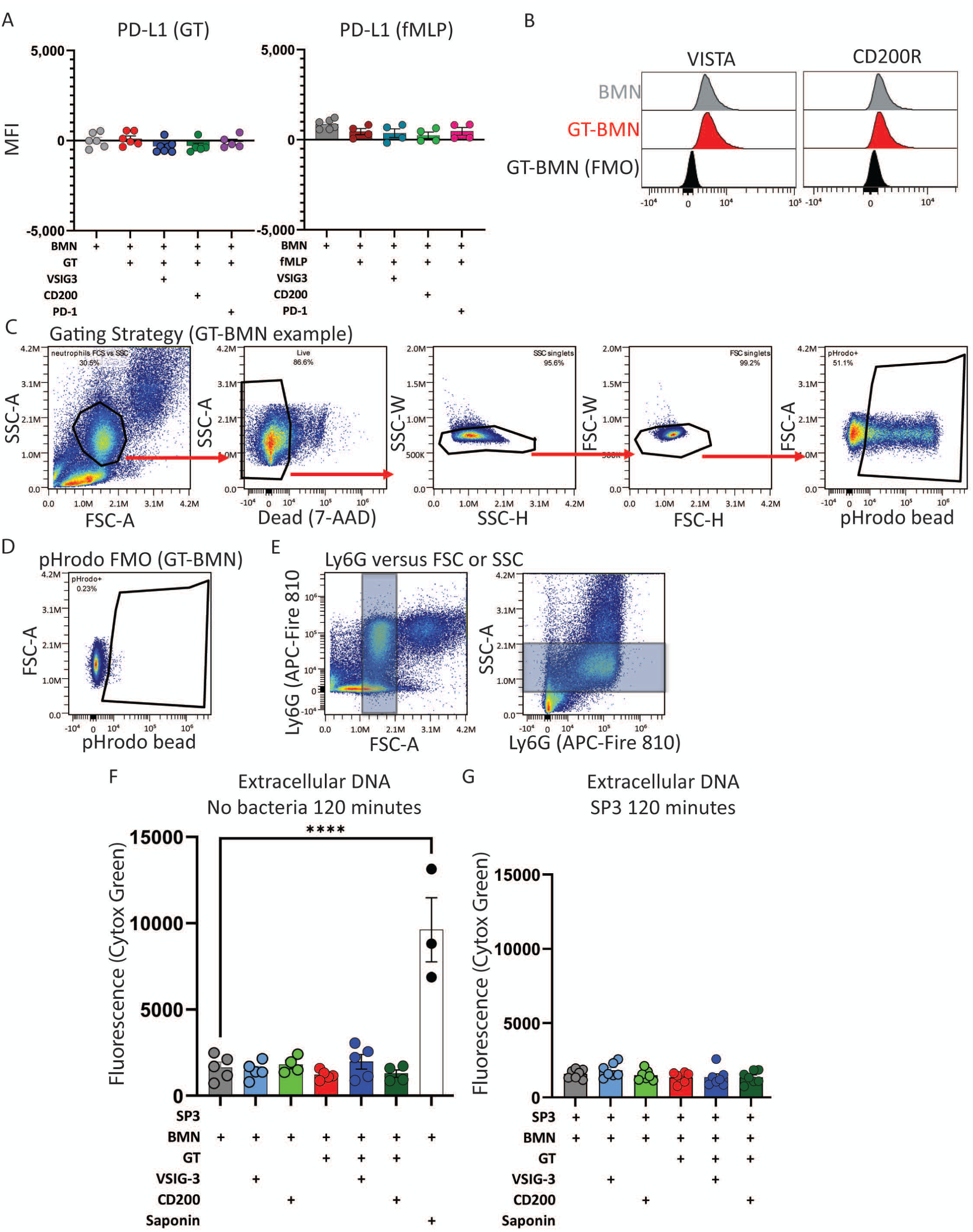
BMN stimulated with or without GT, fMLP, VSIG-3, CD200, or PD-1 were analyzed via flow cytometry to confirm the lack of expression of (**A**) PD-Ll. (**B**) VISTA and CD200R staining along with antibody cocktail that does not contain VISTA or CD200R antibodies (FMO) confirms BMN expression of VISTA and CD200R. (**C**) Example gating strategy of a GT-BMN stimulated sample to isolate neutrophils from FSC, SSC, and 7-AAD without antibody staining. (**D**) pHrodo bead FMO was used to set the pHrodo positive gate, (**E**) and a Ly6G stained sample shows how FSC and SSC gate was determined for Ly6G+ events. Stimulated BMN were stained with Cytox Green to quantify extracellular DNA after 120 minutes co-cultured with (**F**) no bacteria or (**G**) SP3 for 120 minutes. Statistics in bar graphs used were one-way ANOVAs with Bonferroni multiple comparisons corrections or simple linear regression where * = P < 0.05, ** = P < 0.01, *** = P < 0.001, and ****= P < 0.0001. Error bars show standard error of the mean (SEM).

**Supplemental Figure 4.**
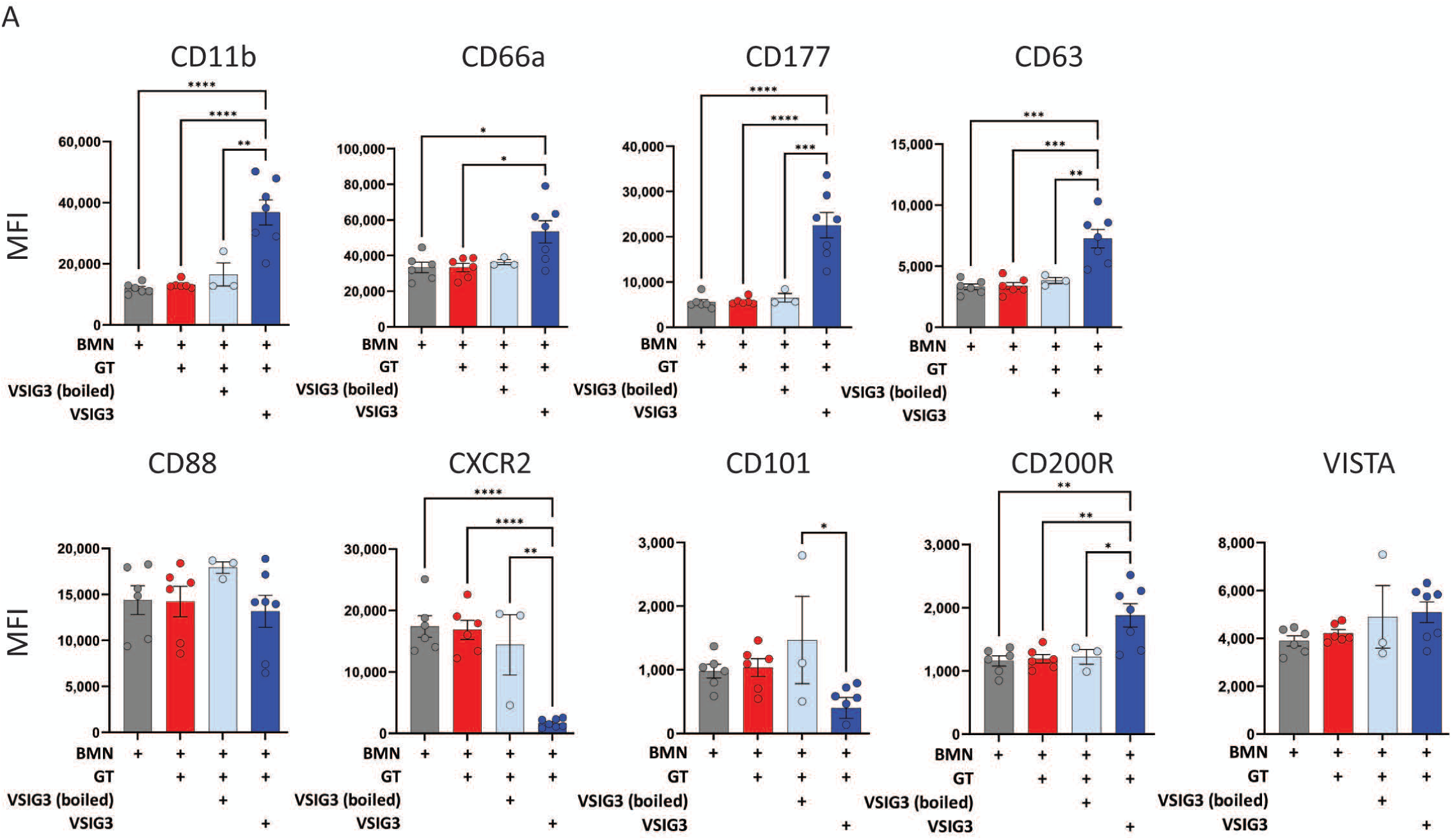
(**A**) BMNs were stimulated with or without GT, heat inactivated (boiled) VSIG-3, or VSIG-3 to validate ligand specific changes in expression. Statistics used were one-way ANOVAs with Bonferroni multiple comparisons corrections where *= P < 0.05, ** = P < 0.01, *** = P < 0.001, and ****= P < 0.0001. Error bars show standard error of the mean (SEM).

